# Pathogenic LRRK2 causes age-dependent and region-specific deficits in ciliation, innervation and viability of cholinergic neurons

**DOI:** 10.1101/2024.07.16.603799

**Authors:** Besma Brahmia, Yahaira Naaldijk, Pallabi Sarkar, Loukia Parisiadou, Sabine Hilfiker

**Affiliations:** Dept. of Anesthesiology, Rutgers New Jersey Medical School, Newark, NJ, USA; Dept. of Physiology, Pharmacology and Neuroscience, Rutgers New Jersey Medical School, Newark, NJ, USA; Dept. of Pharmacology, Northwestern University, Chicago, IL, USA

**Keywords:** Parkinson’s disease, LRRK2, cilia, cholinergic neurons, Rab12

## Abstract

Pathogenic activating point mutations in the LRRK2 kinase cause autosomal-dominant familial Parkinsońs disease (PD). In cultured cells, mutant LRRK2 causes a deficit in *de novo* cilia formation and also impairs ciliary stability. In brain, previous studies have shown that in PD patients due to the G2019S-LRRK2 mutation as well as in middle-aged G2019S-LRRK2 knockin mice, striatal cholinergic interneurons show a deficit in primary cilia. Here, we show that cilia loss in G2019S-LRRK2 knockin mice is not limited to cholinergic striatal interneurons but common to cholinergic neurons across distinct brain nuclei. The lack of cilia in cholinergic forebrain neurons is accompanied by the accumulation of LRRK2-phosphorylated Rab12 GTPase and correlates with the presence of dystrophic cholinergic axons. Those deficits are already evident in young adult mutant LRRK2 mice. In contrast, the age-dependent loss of cilia in brainstem cholinergic neurons correlates with an age-dependent loss of cholinergic innervation derived from this brain area. Strikingly, we find cholinergic cell loss in mutant LRRK2 mice that is age-dependent, cell type-specific and disease-relevant. The age-dependent loss of a subset of cholinergic neurons mimics that observed in sporadic PD patients, highlighting the possibility that these particular neurons may require functional cilia for long-term cell survival.

## Introduction

Autosomal-dominant point mutations in the LRRK2 gene cause late-onset familial Parkinsońs disease (PD) and variants at the LRRK2 locus increase the risk for sporadic PD (Blauwendraat et al., 2020; Nalls et al., 2019). Thus, LRRK2 is key to our understanding of PD pathomechanisms. G2019S-LRRK2 PD patients and non-manifesting mutation carriers show increased brain cholinergic activity, which may represent an early adaptation to compensate for LRRK2-related cholinergic dysfunction (Batzu et al., 2023; S.-Y. Liu et al., 2018). Sporadic PD patients also show cholinergic defects in distinct brain areas that are refractive to dopaminergic therapy and correlate with symptoms typical for the prodromal and early stages of PD (Bohnen et al., 2022; Perez-Lloret and Barrantes, 2016). Neuronal vulnerability in PD is not limited to dopaminergic neurons but has also been observed in specific cholinergic brain nuclei (Giguère et al., 2018; Surmeier et al., 2017). The G2019S-LRRK2 mutation shows incomplete penetrance, suggesting that additional factors contribute to PD in LRRK2 mutation carriers (Healy et al., 2008). These features of human LRRK2-PD are recapitulated in the preclinical G2019S-LRRK2 knock-in (KI) mouse model that expresses pathogenic LRRK2 at physiologically relevant levels and patterns (Yue et al., 2015). G2019S-LRRK2-KI mice show early cholinergic alterations (Hussein et al., 2022). They do not display overt dopaminergic neuronal cell loss or motor phenotypes, but greater susceptibility to dopaminergic cell death induced by a mitochondrial toxin as compared to wildtype mice (Novello et al., 2022; Yue et al., 2015). These findings indicate that G2019S-LRRK2-KI mice are a valuable model for studying early disease-related impairments and the selective cellular vulnerabilities associated with LRRK2-PD.

LRRK2 encodes a protein kinase and pathogenic mutations including the most common G2019S mutation increase its kinase activity (Greggio et al., 2006; Steger et al., 2016; West et al., 2005). A subset of Rab GTPases that are regulators of membrane trafficking serve as kinase substrates (Steger et al., 2017, 2016; Thirstrup et al., 2017). Phosphorylation of Rab proteins interferes with their ability to interact with regulatory and effector proteins (Steger et al., 2017, 2016) that can interfere with their normal function in membrane trafficking (Mamais et al., 2024; Rivero-Ríos et al., 2020, 2019). Importantly, once phosphorylated, the Rab proteins gain the ability to interact with a new set of effector proteins. For Rab8, Rab10 and Rab12 this includes the RILPL1 and RILPL2 proteins (Ito et al., 2023; Steger et al., 2017; Waschbüsch and Khan, 2020). In cultured cells, a centriolar phospho-Rab/RILPL1 protein complex interferes with *de novo* cilia formation and with the process that keeps the two centrioles close to each other, known as centriole cohesion (Dhekne et al., 2018; Fdez et al., 2022; Fernández et al., 2019; Lara Ordónez et al., 2019; Lara Ordóñez et al., 2022; Naaldijk et al., 2024). If formed, the stability of cilia is also negatively influenced by LRRK2 in a manner dependent on the kinase activity but independent of a phospho-Rab/RILPL1 complex (Sobu et al., 2021).

Ciliation defects have been found in striatal cholinergic interneurons in human postmortem brains from G2019S-LRRK2-PD and sporadic PD patients (Khan et al., 2024), as well as in multiple LRRK2 mouse models including in 13-months old G2019S-LRRK2-KI mice (Khan et al., 2021). No ciliary defects have been observed in medium spiny neurons that make up around 95% of neurons in the striatum. This lack of primary cilia in striatal interneurons is the most profound anatomical cellular deficit associated with pathogenic LRRK2 in the intact brain. However, it remains unknown whether pathogenic LRRK2 affects ciliation in other cholinergic cell types, and how a lack of cilia may impact on cholinergic innervation and/or the viability of cholinergic neurons as relevant for prodromal and early PD. Here, we report age-dependent, region-specific and cell type-specific ciliation defects in distinct cholinergic nuclei in G2019S-LRRK2-KI mice. The differential temporal pattern of these changes across brain areas implies LRRK2-mediated effects on both cilia formation and ciliary stability. Notably, we find that ciliation defects in cholinergic neurons correlate with deficient innervation and the age-dependent death of a subset of cholinergic neurons that are also vulnerable in human PD.

## Results

### Age-dependent primary cilia defects in basal ganglia cholinergic neurons of G2019S-LRRK2-KI mice

Cholinergic interneurons in the dorsal striatum of G2019S-LRRK2 knock-in (KI) mice demonstrate a ciliary deficit at 13 months of age (Khan et al., 2021). Since age is a primary risk factor for PD (Domingo and Klein, 2018), we chose to measure ciliation in different neuronal cell populations across ages to determine whether ciliary deficits occur with age. Because postnatal maturation of primary cilia in mouse brain takes 8-12 weeks (Arellano et al., 2012; Brewer et al., 2024), we chose our earliest timepoint for cilia analysis in young adult mice (4-5 months). As comparison, we used middle-aged (13-14 months) mice that correlate with human ages in the late-forties, and aged mice (18-24 months) that correlate with human ages in the late fifties and sixties (Jackson et al., 2017). We evaluated ciliation in choline acetyltransferase-positive (ChaT+) striatal interneurons in the caudate putamen (CPu) at different timepoints and find that there is an age-dependent loss of cilia in G2019S-LRRK2-KI as compared to wildtype (wt) mice which is present in both males and females (**Figure 1A-D**). The ciliary deficit is not apparent until mice reach middle-age, and there is no further loss of cilia in aged mice (**Figure 1B-D**). We separately determined ciliation in the dorsomedial and ventrolateral CPu since cholinergic neurons in those two striatal compartments show distinct intrinsic baseline activities (Ahmed et al., 2019), but we did not detect differences in their susceptibility to ciliary loss. In the dorsomedial striatal cholinergic neurons which remain ciliated in mutant aged mice, we saw a slight increase in ciliary length that may reflect an increase in ciliary signaling capacity (Guo et al., 2017) (**Figure 1E**).

**Figure 1.**
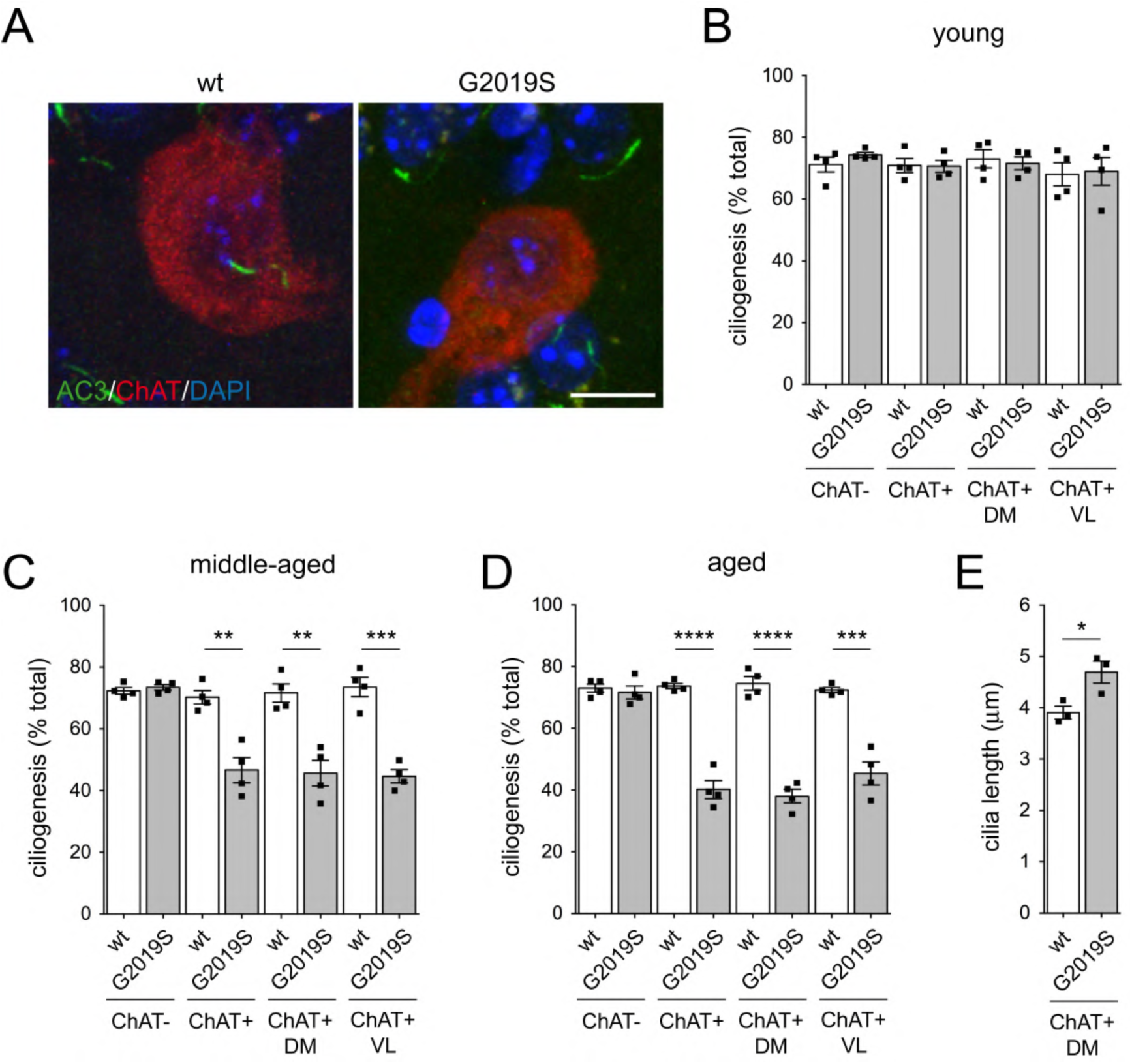
Age-dependent loss of primary cilia in cholinergic interneurons in caudate putamen (CPu) of G2019S-LRRK2-KI mice. (**A**) Confocal images of cholinergic interneurons from aged wildtype (wt) and G2019S-LRRK2-KI (G2019S) mice stained for neuronal ciliary marker adenylate cyclase 3 (AC3), choline acetyltransferase (ChaT) and DAPI. Scale bar, 10 μm. (**B**) Quantitation of ciliation in ChaT+ and ChaT-neurons in CPu from young (4-5 months) wt and G2019S mice. Ciliation in ChaT+ neurons was also quantified separately from dorsomedial (DM) and ventrolateral (VL) CPu. (**C**) Ciliation in ChaT+ and ChaT-neurons in middle-aged (13-14 months) wt and G2019S mice. (**D**) Ciliation in aged (18-24 months) mice. Each datapoint is from an individual mouse, analyzing 2 sections per animal and 100-200 ChaT+ and 400-800 ChaT-cells. Bars represent mean ± s.e.m (n=4, 2 males and 2 females per age group and genotype). Significance was determined by unpaired two-tailed student’s T-test; **p<0.01; ***p<0.001; ****p<0.0001. (**E**) Quantitation of ciliary length in ChaT+ neurons from aged (18-24 months old) wt and G2019S mice. Bars represent mean ± s.e.m (n=3); *p<0.05.

To determine whether ciliary defects were exclusive to cholinergic interneurons, we measured ciliation in two subsets of GABAergic interneurons (Tepper et al., 2010) in CPu from aged animals. Parvalbumin-positive (PV+) and calretinin-positive (calret+) neurons showed no difference in ciliation in G2019S-LRRK2-KI mice (**Figure 1 – figure supplement 1**). We also measured ciliation in non-ChaT+ neurons, which we assume are mainly comprised of medium spiny neurons that make up the vast majority of striatal neurons. Consistent with previous reports (Dhekne et al., 2018; Khan et al., 2021), no differences in ciliation were observed in ChaT-cells between wt and mutant LRRK2 mice across all age groups (**Figure 1 B-D**).

We next wondered whether the age-dependent ciliary defect was unique to striatal cholinergic interneurons. The globus pallidus (GP) is another basal ganglia nucleus that contains cholinergic neurons, and we found a similar age-dependent ciliation defect as in the dorsal striatum, with a significant loss of cilia observed only in middle-aged and aged mutant mice (**Figure 2**). When measuring ciliated nuclei of ChaT-cells in the GP, we did not see any change in percent ciliation across all age groups. Together, these data show an age-dependent ciliary defect in cholinergic neurons in the striatum and GP from G2019S-LRRK2-KI mice.

**Figure 2.**
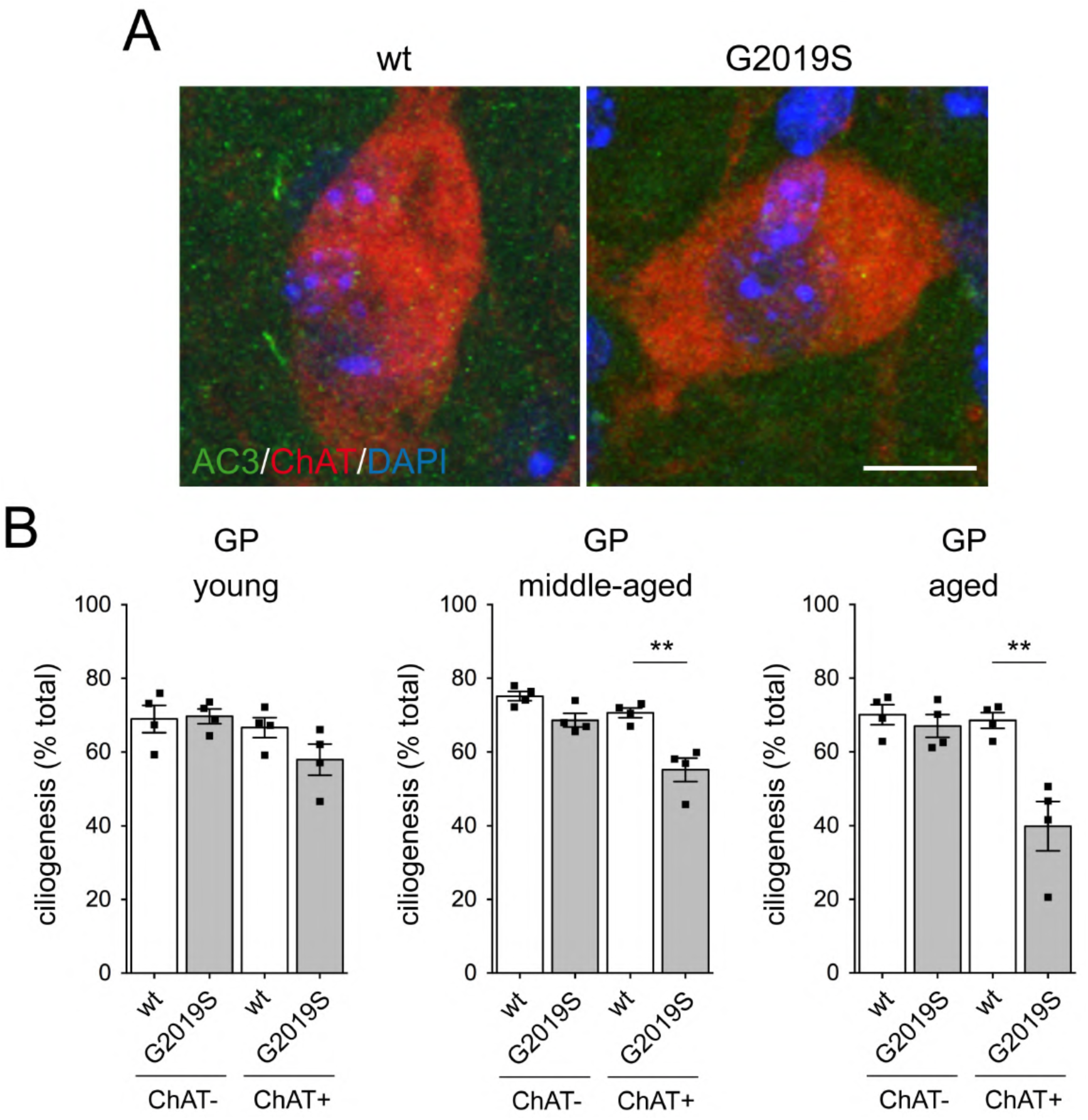
Age-dependent loss of primary cilia in cholinergic neurons in globus pallidus (GP) of G2019S-LRRK2-KI mice. (**A**) Confocal images of cholinergic neurons in GP from aged wildtype (wt) and G2019S-LRRK2-KI (G2019S) mice stained for neuronal ciliary marker adenylate cyclase 3 (AC3), choline acetyltransferase (ChaT) and DAPI. Scale bar, 10 μm. (**B**) Quantitation of ciliation in ChaT+ and ChaT-neurons in CPu from young (4-5 months), middle-aged (13-14 months) and aged (18-24 months) wt and G2019S mice. Each datapoint is from an individual mouse, analyzing 2 sections per animal and 100-200 ChaT+ and 400-800 ChaT-cells. Bars represent mean ± s.e.m (n=4, 2 males and 2 females per age group and genotype). Significance was determined by unpaired two-tailed student’s T-test; **p<0.01.

### Ciliary defects in basal forebrain and brainstem cholinergic neurons in G2019S-LRRK2-KI mice

Cholinergic neurons in the basal forebrain are a main source of acetylcholine in the central nervous system and widely project to many brain areas. They are classified into four distinct subpopulations located in the medial septum (MS), the vertical and horizontal limbs of the diagonal band of Broca (VDB/HDB), and the nucleus basalis of Meynert (NBM) (Ananth et al., 2023).We measured ciliation in ChaT+ neurons in the NBM and notably found a ciliary deficit of the same magnitude as that observed in striatal cholinergic neurons already in young mice that persisted in middle-aged and aged mice (**Figure 3A,B**). Similarly, basal forebrain cholinergic neurons in the MS and DB showed an early onset of the ciliary deficit, while there was no change in percent ciliation of ChAT-cells in any of the regions analyzed (**Figure 3C,D)**.

**Figure 3.**
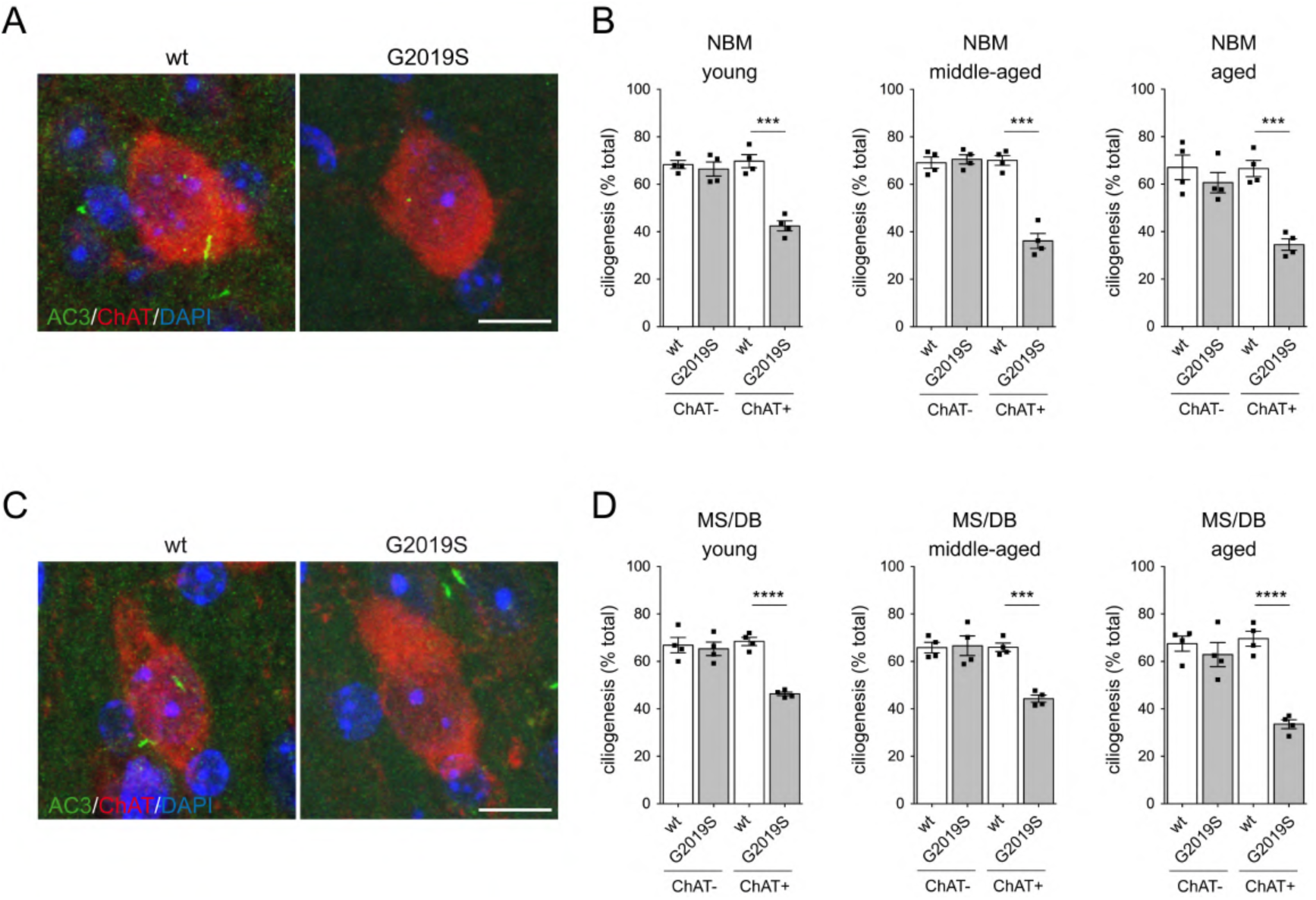
Ciliary deficits in forebrain cholinergic neurons already in young G2019S-LRRK2-KI mice. (**A**) Confocal images of cholinergic neurons in nucleus basalis of Meynert (NBM) from aged wt and G2019S mice stained for neuronal ciliary marker adenylate cyclase 3 (AC3), choline acetyltransferase (ChaT) and DAPI. Scale bar, 10 μm. (**B**) Quantitation of ciliation in ChaT+ and ChaT-neurons in NBM from young (4-5 months), middle-aged (13-14 months) and aged (18-24 months) wt and G2019S mice. (**C**) Confocal images of cholinergic neurons in medial septum (MS) from aged wt and G2019S mice stained for neuronal ciliary marker adenylate cyclase 3 (AC3), choline acetyltransferase (ChaT) and DAPI. Scale bar, 10 μm. (**D**) Quantitation of ciliation in ChaT+ and ChaT-neurons in MS/diagonal band (DB) from young (4-5 months), middle-aged (13-14 months) and aged (18-24 months) wt and G2019S mice. Each datapoint is from an individual mouse, analyzing 2 sections per animal and 100-200 ChaT+ and 400-800 ChaT-cells. Bars represent mean ± s.e.m (n=4, 2 males and 2 females per age group and genotype). Significance was determined by unpaired two-tailed student’s T-test; ***p<0.001; ****p<0.0001.

Cholinergic projection neurons are also found in two brainstem nuclei, the pedunculopontine nucleus (PPN) and the laterodorsal tegmental nucleus (LDTgN). Using the same approach to evaluate ciliation, we found an age-dependent ciliary deficit in cholinergic neurons in both the PPN and LDTgN that was evident in middle-aged LRRK2-G2019S-KI mice and did not worsen with age (**Figure 4**). Together, these data show that G2019S-LRRK2-KI mice have primary cilia defects in all cholinergic neuronal cell populations analyzed here, but with differences in age-dependent onset. Cholinergic basal forebrain neurons show a ciliation defect already in young adult mice, whilst basal ganglia and brainstem cholinergic neurons lose their cilia with age.

**Figure 4.**
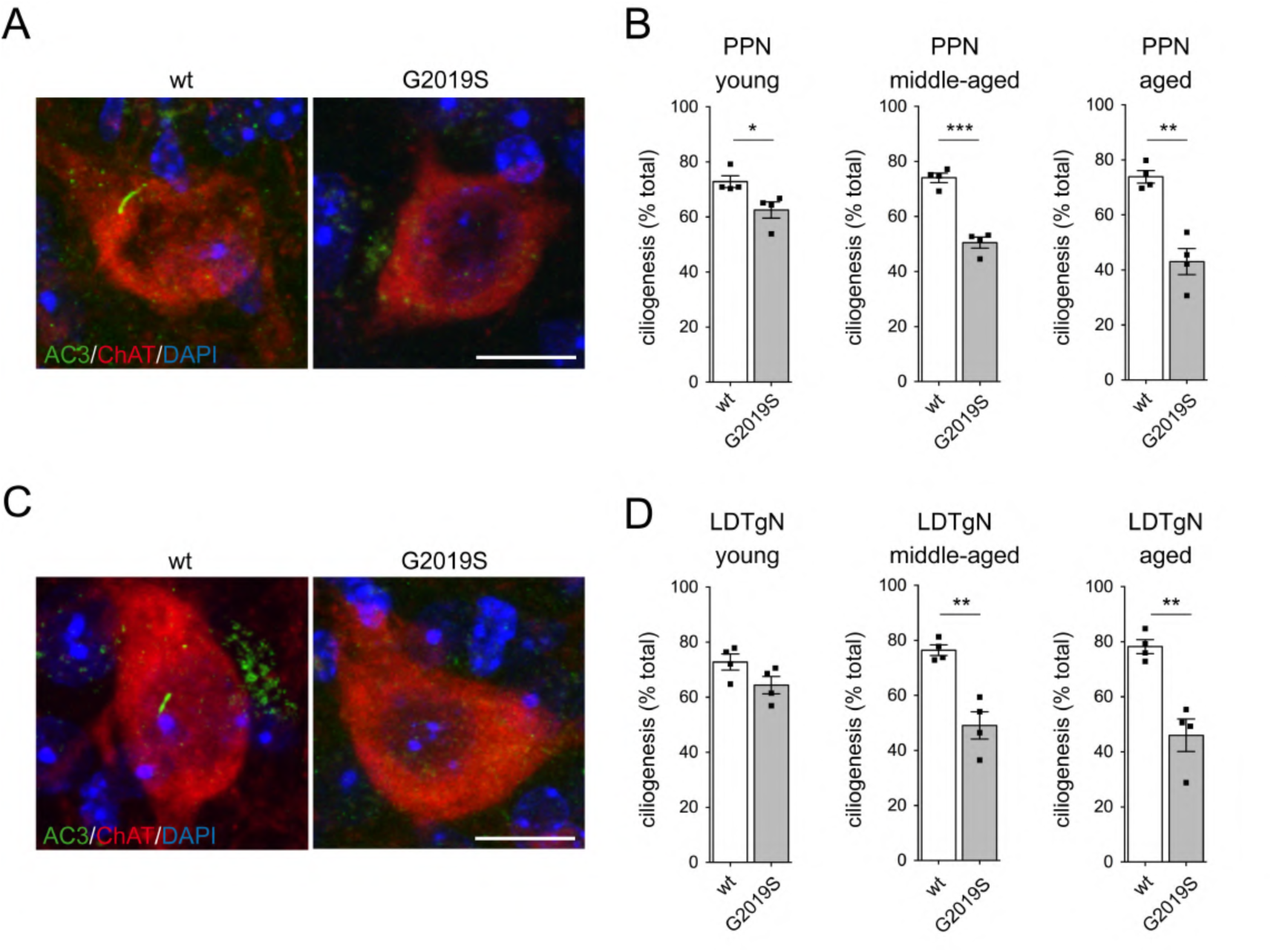
Age-dependent loss of primary cilia in brainstem cholinergic neurons in G2019S-LRRK2-KI mice. (**A**) Confocal images of cholinergic neurons in pedunculopontine nucleus (PPN) from aged wt and G2019S mice stained for neuronal ciliary marker adenylate cyclase 3 (AC3), choline acetyltransferase (ChaT) and DAPI. Scale bar, 10 μm. (**B**) Quantitation of ciliation in ChaT+ neurons in PPN from young (4-5 months), middle-aged (13-14 months) and aged (18-24 months) wt and G2019S mice. (**C**) Confocal images of cholinergic neurons in laterodorsal tegmental nucleus (LDTgN) from aged wt and G2019S mice stained for neuronal ciliary marker adenylate cyclase 3 (AC3), choline acetyltransferase (ChaT) and DAPI. Scale bar, 10 μm. (**D**) Quantitation of ciliation in ChaT+ neurons in LDTgN from young (4-5 months), middle-aged (13-14 months) and aged (18-24 months) wt and G2019S mice. Each datapoint is from an individual mouse, analyzing 2 sections per animal and 100-200 ChaT+ cells. Bars represent mean ± s.e.m (n=4, 2 males and 2 females per age group and genotype). Significance was determined by unpaired two-tailed student’s T-test; *p<0.05; **p<0.01; ***p<0.001.

### Phosphorylated Rab12 in basal forebrain cholinergic neurons

Previous *in vitro* work by us and others shows the importance of LRRK2-phosphorylated Rab10 and its effector, RILPL1 in blocking primary cilia formation (Dhekne et al., 2018; Lara Ordónez et al., 2019). Rab12 promotes LRRK2 activation both basally and in response to lysosomal damage, which leads to an increase in lysosomal phospho-Rab10 (pRab10) and phospho-Rab12 (pRab12) (Dhekne et al., 2023; Wang et al., 2023), and both pRab10 and pRab12 interact with RILPL1 (Ito et al., 2023; Steger et al., 2017). We therefore wondered whether the ciliation defects in cholinergic neurons mediated by G2019S-LRRK2 correlated with the appearance of phosphorylated Rab substrates. To explore the possible contribution of pRab10 and pRab12 to ciliogenesis blockade in cholinergic neurons, we employed available phospho-state-specific antibodies. Both pRab10 and pRab12 antibodies are Rab-specific and phospho-state-specific as assessed by western blotting (Kalogeropulou et al., 2020; Mir et al., 2018). In our hands and as previously described (Khan et al., 2021), the pRab10 antibody did not stain mouse brain tissue sections. In contrast, the pRab12 antibody revealed brightly stained pRab12+ punctae in basal forebrain cholinergic neurons from G2019S-LRRK2-KI as compared to control mice (**Figure 5A**). The prevalence of these pRab12+ structures as well as the intensity of cytosolic pRab12 staining seemed to worsen with age, so we chose to quantify both the number of pRab12+ structures as well as integrated density as a measure of pRab12 staining intensity from the same neurons. We found a significant increase in the number of pRab12+ structures in cholinergic neurons in the HDB and MS, and a significant increase in pRab12 staining intensity in cholinergic neurons in the HDB and MS, in 13+ months old G2019S-LRRK2-KI mice as compared to controls (**Figure 5B,C**). In contrast to basal forebrain cholinergic neurons, no pRab12+ punctae were observed in cholinergic neurons in brainstem, CPu or GP from G2019S-LRRK2-KI mice (**Figure 5 – figure supplement 1**). To confirm that the pRab12 antibody was detecting a phospho-specific epitope, we subjected slices to phosphatase treatment before staining, which completely abolished both punctate and cytosolic pRab12 staining (**Figure 5 – figure supplement 2**). In addition, and using an antigen retrieval protocol, we found that the punctate pRab12 staining looked vesicular in nature (**Figure 5 – figure supplement 2**). Altogether, these data show that the ciliogenesis defect in basal forebrain cholinergic neurons from G2019S-LRRK2-KI mice correlates with a vesicular accumulation of pRab12 (or a phospho-epitope detected by the pRab12 antibody) that is evident already in young adult mice, worsens with age, and is not present in other cholinergic neuronal cell populations.

**Figure 5.**
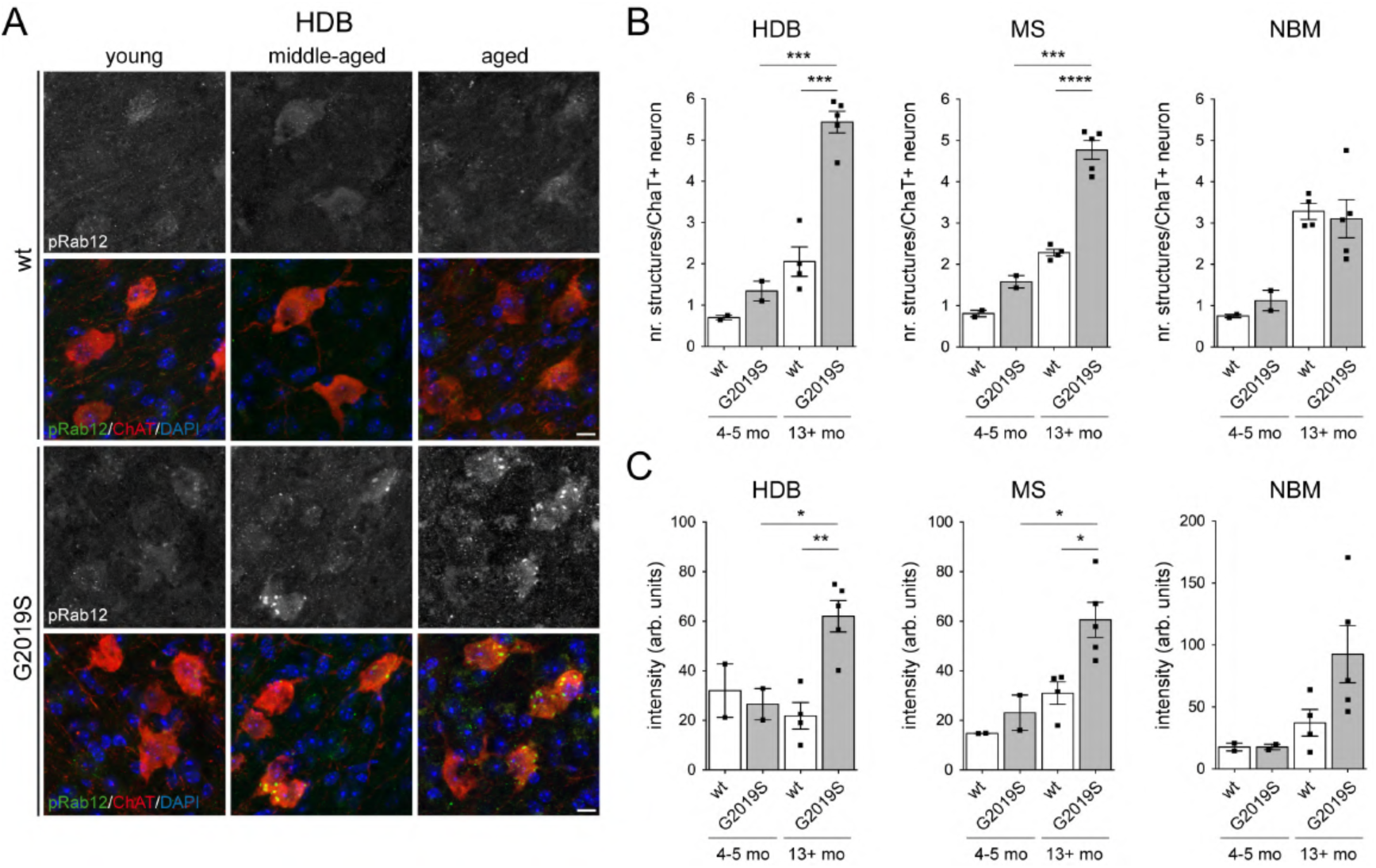
Basal forebrain cholinergic neurons from G2019S-LRRK2-KI mice show unique pRab12 staining. (**A**) Confocal images of cholinergic neurons in horizontal diagonal band (HDB) from young, middle-aged and aged wt and G2019S mice stained with antibodies against phospho-Rab12 (pRab12), choline acetyltransferase (ChaT) and DAPI. Scale bar, 10 μm. (**B**) Quantification of pRab12-positive structures in ChaT+ neurons in HDB, MS and NBM from young adult and 13+ months old wt and G2019S mice. Data are from young adult (n=2) and 13+ months old wt (n=4) and G2019S (n=5) mice, with 50-100 ChaT+ neurons from 2 independent sections quantified per animal. (**C**) Quantification of pRab12 fluorescence intensity in ChaT+ neurons in HDB, MS and NBM from young adult and 13+ months old wt and G2019S mice. Integrated density from young adult (n=2) amd 13+ months old wt (n=4) and G2019S (n=5) mice, with 20-45 ChaT+ neurons from 2 independent sections quantified per animal. Significance was determined by unpaired two-tailed student’s T-test; ***p<0.001; ****p<0.0001.

### Distinct deficits in innervation derived from forebrain and brainstem cholinergic neurons in G2019S-LRRK2-KI mice

The vesicular pRab12 staining in basal forebrain cholinergic neurons is consistent with an early defect in *de novo* cilia formation (Dhekne et al., 2018; Lara Ordónez et al., 2019). Since a lack of cilia can lead to axonal growth defects (Guadiana et al., 2013; Guo et al., 2019), we next characterized cholinergic innervation. Cholinergic neurons project across many domains, but certain target regions receive preferential innervation from distinct cholinergic nuclei, including the dorsal hippocampal CA1/CA2 regions, where projections are mainly derived from cholinergic neurons in the MS/DB (Li et al., 2018). Using vesicular acetylcholine transporter (VAChT) labeling, we found a similar reticular pattern of immunoreactivity in the dorsal hippocampus of G2019S-LRRK2-KI and control mice, reflecting a dense network of cholinergic axons. However, in addition, we observed sparse, large axonal clusters that were already present in young adult G2019S-LRRK2 mice (**Figure 6A**). These clusters represented abnormal cholinergic axons as confirmed by double staining with ChaT (**Figure 6 – figure supplement 1**). When quantifying the number of dystrophic axonal clusters and the total number of dystrophic axons, we found an increase in young G2019S-LRRK2-KI mice as compared to control mice, and a worsening of the dystrophic axonal phenotype with age in the mutant mice (**Figure 6B,C**).

**Figure 6.**
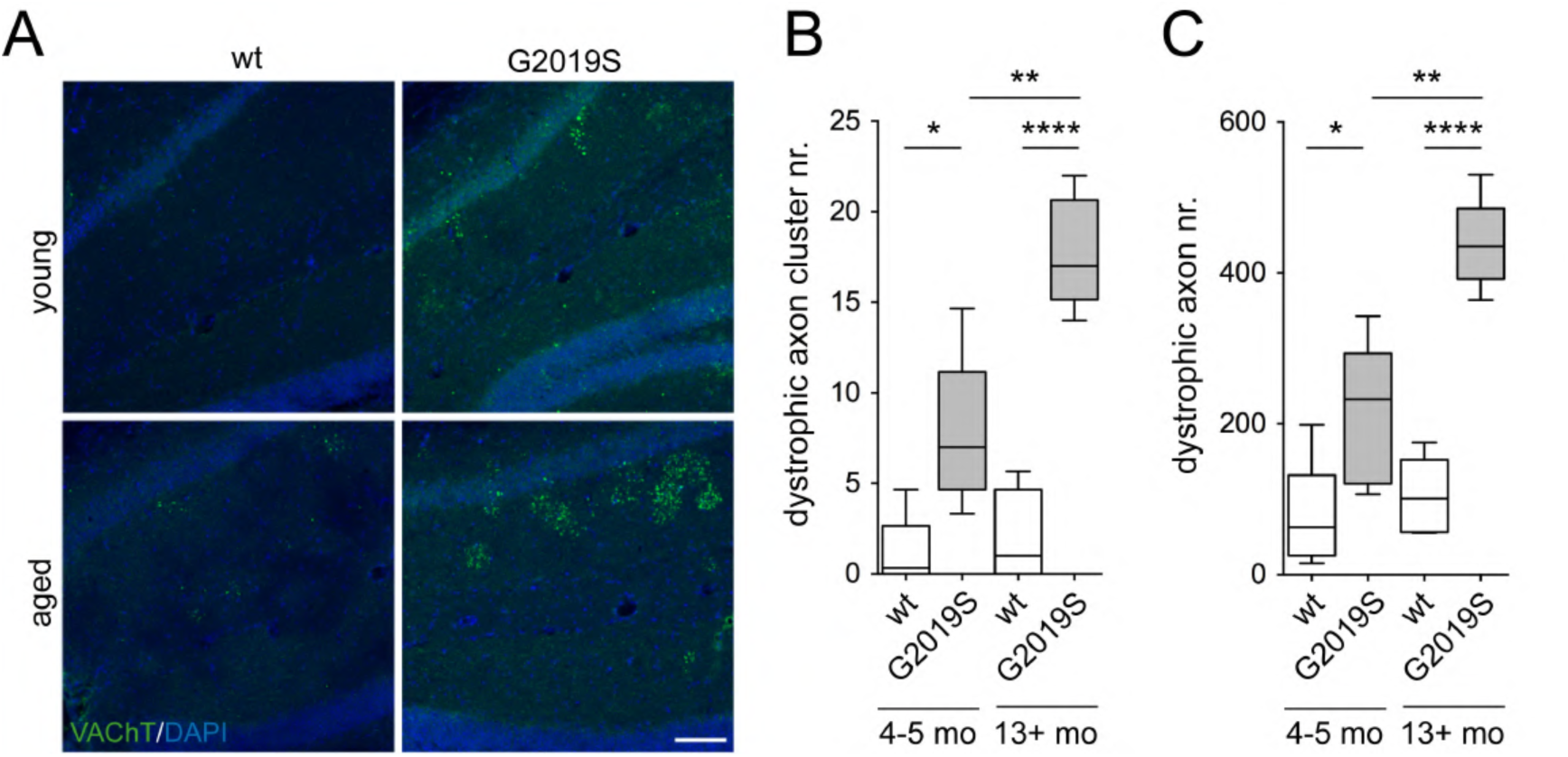
Early dystrophic changes in cholinergic axons derived from basal forebrain cholinergic neurons in G2019S-LRRK2-KI mice. (**A**) Confocal images of dorsal hippocampal CA1/2 regions from young and aged wt and G2019S mice stained for vesicular acetylcholine transporter (VAChT) and DAPI. Scale bar, 100 μm. (**B**) Manual quantitation of dystrophic axonal clusters in hippocampal CA1/CA2 regions from young (4-5 months) and 13+ months old wt and G2019S mice. (**C**) Quantitation of dystrophic axons > 3 μm^2^ in hippocampal CA1/CA2 regions from young (4-5 months) and 13+ months old wt and G2019S mice. Box and whisker plots (5-95 percentile) are from 5 animals (male and female) each per age and genotype, with three sections per animal and three images per section quantified. Significance was determined by unpaired two-tailed student’s T-test; *p<0.05; **p<0.01; ****p<0.0001.

We next wondered whether the age-dependent ciliary loss in cholinergic brainstem neurons correlates with age-dependent deficits in innervation. As a target area for analysis, we chose the thalamic dorsal lateral geniculate nucleus (dLGN), as this region receives preferential innervation from cholinergic neurons in the PPN (Huerta-Ocampo et al., 2020). Cholinergic innervation of the dLGN was not altered in young adult mice, but a deficit was evident in middle-aged and aged mutant LRRK2 mice (**Figure 7**). As above, we performed co-staining with ChaT to confirm that we were quantifying cholinergic axons (**Figure 7 – Figure supplement 1**). Altogether, our data show that dystrophic changes in axons derived from basal forebrain cholinergic neurons are already present in young adult G2019S-LRRK2-KI mice, correlate with the early ciliary defects, and worsen in an age-dependent manner. In contrast, brainstem cholinergic neurons show a deficit in innervation only with age that correlates with the age-dependent loss of cilia.

**Figure 7.**
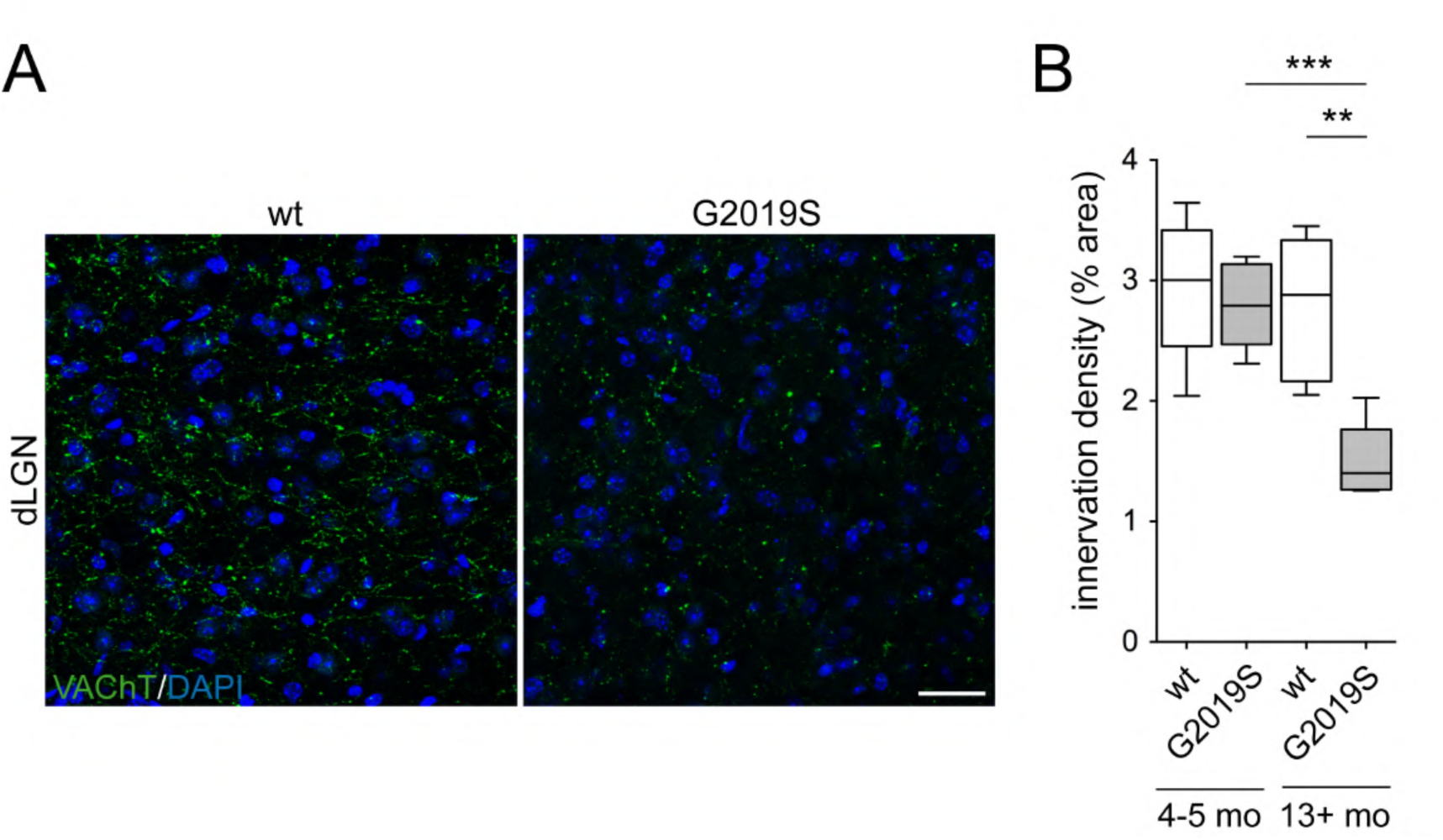
Age-dependent loss of cholinergic innervation derived from brainstem cholinergic neurons in G2019S-LRRK2-KI mice. (**A**) Confocal images of thalamic dorsal lateral geniculate nucleus (dLGN) from aged wt and G2019S mice stained for vesicular acetylcholine transporter (VAChT) and DAPI. Scale bar, 30 μm. (**B**) Quantitation of innervation density in dLGN from young (4-5 months) and 13+ months old wt and G2019S mice. Box and whisker plots (5-95 percentile) are from 5 animals (male and female) each per age and genotype, with three sections per animal and five images per section quantified. Significance was determined by unpaired two-tailed student’s T-test; **p<0.01; ***p<0.001.

### Age-dependent cholinergic cell loss in select forebrain and brainstem nuclei in G2019S-LRRK2-KI mice

Whilst most neurons in the adult brain are ciliated (Arellano et al., 2012), the role of cilia for neuronal integrity and viability remains largely unknown. To determine whether a lack of cilia may affect ChaT+ cell survival, we performed bilateral cell counts (Chen et al., 2020; González-Rodríguez et al., 2021) in cholinergic cells of the basal ganglia, basal forebrain and brainstem from young, middle-aged and aged G2019S-LRRK2-KI and control mice. We found no difference in the number of cholinergic cells in the CPu of aged G2019S-LRRK2-KI mice (**Figure 8A**). Similarly, the number of cholinergic neurons in the GP and in the brainstem LDTgN from aged mutant mice was not different from controls (**Figure 8B,C**). In contrast, we observed a significant decrease in ChaT+ cell counts in the NBM from G2019S-LRRK2-KI mice that was apparent already in middle-aged mice and did not worsen with age (**Figure 8D,E**).

**Figure 8.**
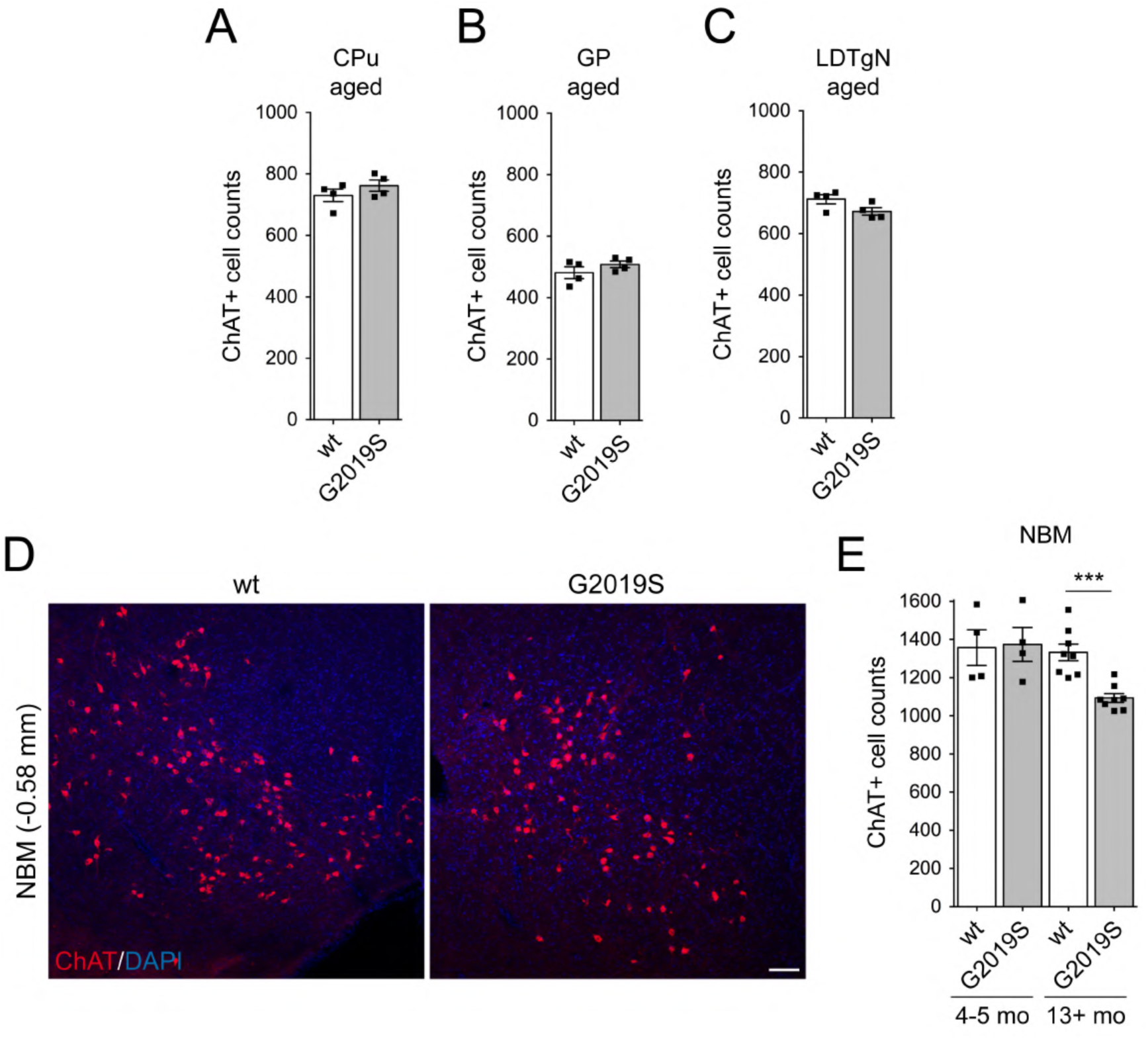
Age-dependent loss of ChaT+ cell counts in nucleus basalis of Meynert in G2019S-LRRK2-KI mice. (**A**) ChaT+ cell counts in CPu from aged wt and G2019S mice. Bars represent mean ± s.e.m (n=4, 2 males and 2 females per genotype). (**B**) ChaT+ cell counts in GP from aged wt and G2019S mice. (**C**) ChaT+ cell counts in brainstem LDTgN from aged wt and G2019S mice. (**D**) Confocal images of NBM from aged wt and G2019S mice stained for ChaT and DAPI, with Bregma coordinates indicated to the left. Scale bar, 100 μm. (**E**) Quantitation of ChaT+ cell counts in NBM from young (4-5 months) and 13+ months old wt and G2019S mice. ChaT+ cells were counted from seven serial 30 μm coronal sections across the NBM. Bars represent mean ± s.e.m (n=4 for young adult, n=8 for 13+ months). Significance was determined by unpaired two-tailed student’s T-test; ***p<0.001.

Since non-cholinergic neurons significantly outnumber cholinergic neurons in the different brain nuclei analyzed, cell counts using NeuN will not show a significant change in the overall number of neurons (Pappas et al., 2018). We therefore co-stained slices from the different cholinergic nuclei with antibodies against either the neurotrophin receptor p75 or neuronal nitric oxide synthase (nNOS) (**Figure 9 – Figure supplements 1,2**). As previously described, we found that all ChaT+ neurons were positive for p75 in basal forebrain MS and DB (Boskovic et al., 2019). Conversely, all ChaT+ neurons were positive for nNOS in brainstem PPN and LDTgN, respectively (Dawson et al., 1991; Geula et al., 1993). This allowed us to employ these markers along with ChaT to determine cholinergic neuronal cell loss as opposed to loss of a cholinergic phenotype. Given the rostro-caudal anatomical distribution of the MS and DB, we serially sectioned and sampled alternating sections that span the MS, VDB and rostral HDB. We found a pronounced decrease in ChaT+ cell counts in the VDB and HDB but not the MS in middle-aged G2019S-LRRK2-KI mice that did not worsen with age (**Figure 9**). An identical decrease was observed when counting p75-positive cells (**Figure 9 – figure supplements 3,4,5**). In contrast, when determining ChaT+/p75+ cell numbers in young adult mice, we found no difference in either the MS, VDB or HDB between G2019S-LRRK2-KI and wt mice (**Figure 9**).

**Figure 9.**
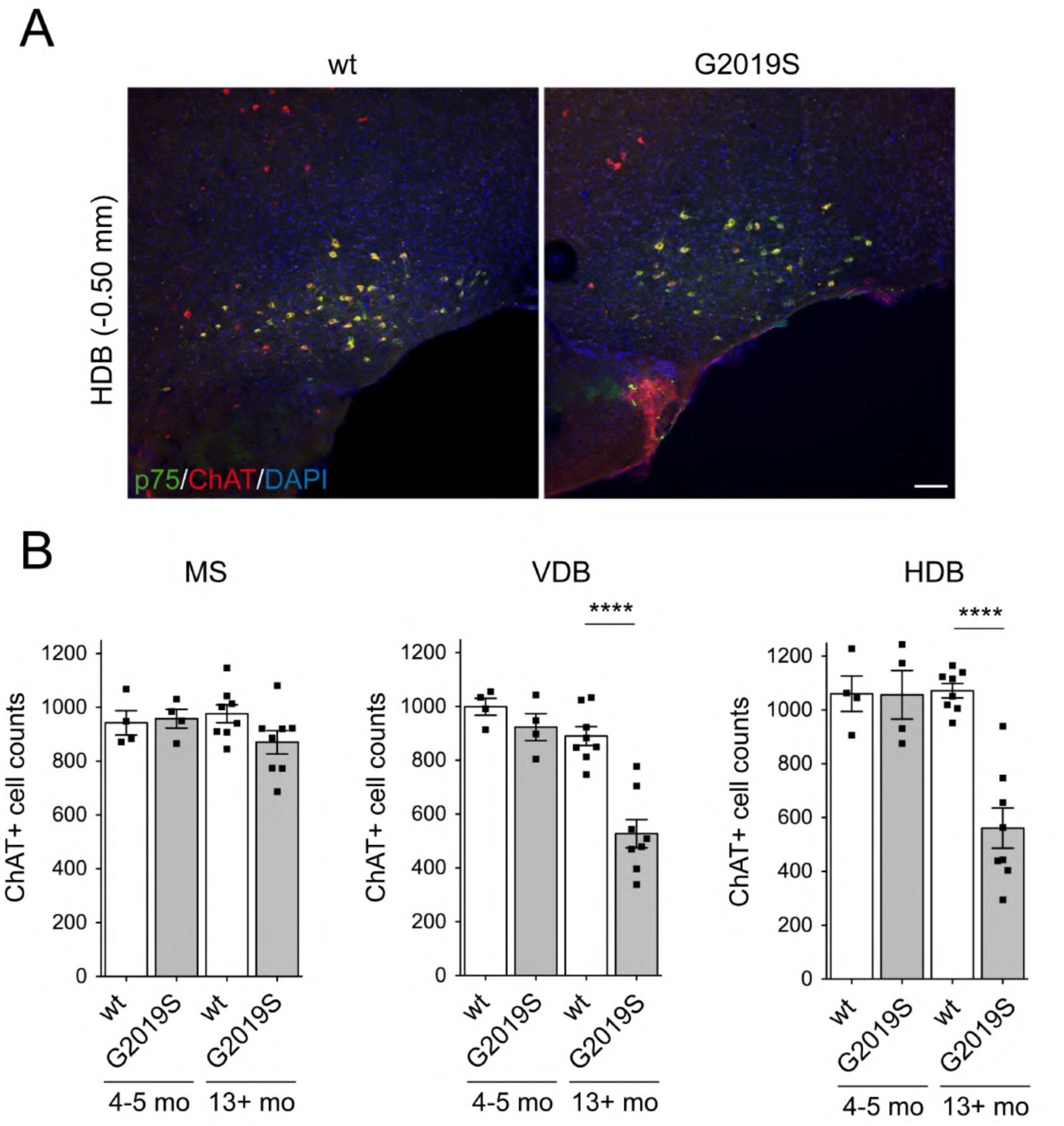
Age-dependent cholinergic cell loss in diagonal band in G2019S-LRRK2-KI mice. (**A**) Confocal images of horizontal diagonal band (HDB) from aged wt and G2019S mice stained for ChaT, p75 and DAPI, with Bregma coordinates indicated to the left. Scale bar, 100 μm. (**B**) Quantitation of ChaT+ cell counts in medial septum (MS), vertical diagonal band (VDB) and horizontal diagonal band (HDB) from young adult (4-5 months) and 13+ months old wt and G2019S mice. ChaT+ cells were counted from fifteen alternate serial 30 μm coronal sections across the MS, VDB and HDB. Bars represent mean ± s.e.m (n=4 for young adult, n=8 for 13+ months). Significance was determined by unpaired two-tailed student’s T-test; ****p<0.0001.

Finally, we quantified cholinergic neurons in the brainstem PPN. Since this nucleus has been divided into two regions based on the density of cholinergic neurons (Martinez-Gonzalez et al., 2011; Mena-Segovia et al., 2009), we serially sectioned and sampled alternating sections for cholinergic neurons from both rostral and caudal PPN (**Figure 10**). A significant loss of cholinergic neurons as assessed by ChaT+/nNOS+ co-staining was observed in caudal but not rostral PPN in middle-aged and aged G2019S-LRRK2-KI mice, whilst we observed no cell loss in young mice (**Figure 10**; **Figure 10 – figure supplement 1**). Therefore, our data for the first time report age-dependent cell death in G2019S-LRRK2-KI mice, which is limited to a subset of cholinergic neurons that also show selective vulnerability in human PD (Giguère et al., 2018; Surmeier et al., 2017).

**Figure 10.**
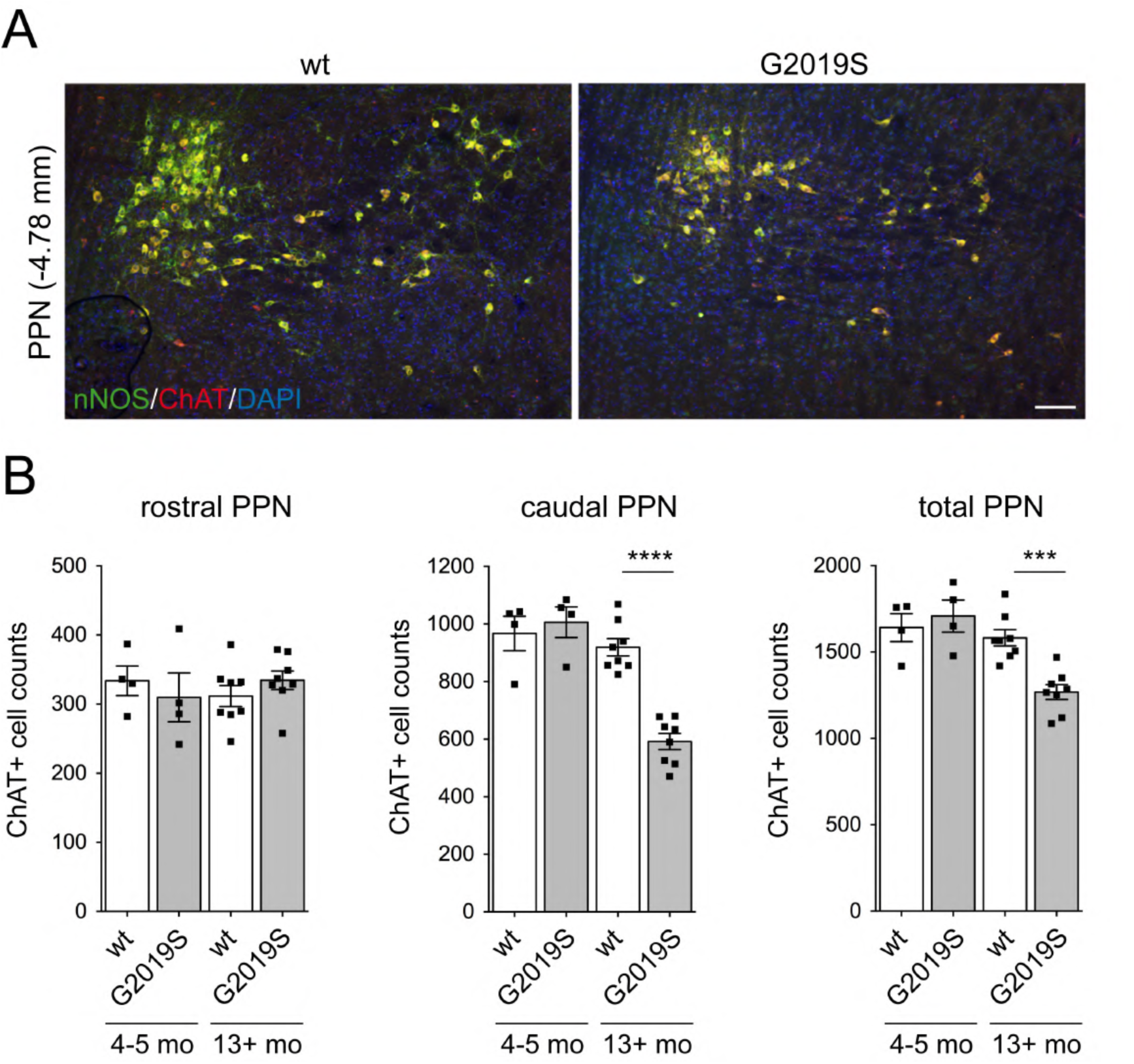
Age-dependent cholinergic cell loss in pedunculopontine nucleus in G2019S-LRRK2-KI mice. (**A**) Confocal images of pedunculopontine nucleus (PPN) from aged wt and G2019S mice stained for ChaT, nNOS and DAPI, with Bregma coordinates indicated to the left. Scale bar, 100 μm. (**B**) Quantitation of ChaT+ cell counts in rostral PPN, caudal PPN and total PPN from young adult (4-5 months) and 13+ months old wt and G2019S mice. ChaT+ cells were counted from fifteen alternate serial 30 μm coronal sections across the PPN (total PPN), as well as from the first six alternate serial rostral sections or the last six alternate serial or caudal sections, respectively. Bars represent mean ± s.e.m (n=4 for young adult, n=8 for 13+ months). Significance was determined by unpaired two-tailed student’s T-test; ***p<0.001; ****p<0.0001.

## Discussion

Here, we find an age-dependent loss of cilia in cholinergic neurons in caudate putamen, globus pallidus and in two brainstem nuclei that is present in middle-aged but not young G2019S-LRRK2-KI mice. In contrast, cholinergic neurons across various forebrain nuclei show a ciliation defect already in young adult mice, which is sustained throughout the animaĺs lifespan. When quantifying ciliation in ChaT-neurons, we did not observe differences in any of the brain areas sampled. Therefore, all cholinergic neurons show a ciliation defect in G2019S-LRRK2-KI mice, but with differences in age-dependent onset. Future work using conditional knockin mice expressing mutant LRRK2 only in cholinergic neurons will be required to determine whether the ciliary defects are due to cell-autonomous mechanism(s).

The age-dependent differences in the onset of the ciliary defects are consistent with the possibility of two distinct underlying mechanisms, one related to an early block in cilia formation and the other related to an age-dependent decrease in ciliary stability. *In vitro* data show that pathogenic LRRK2 causes a block in cilia formation via formation of a pRab/RILPL1 complex (Dhekne et al., 2018, p. 201; Lara Ordónez et al., 2019), and it will be important for us to determine the earliest LRRK2-mediated effects on *de novo* cilia formation in cholinergic forebrain neurons by analyzing ciliation from postnatal and young mutant LRRK2 animals. The lack of cilia in basal forebrain cholinergic neurons from young adult G2019S-LRRK2-KI mice is accompanied by the striking presence of pRab12-positive vesicular structures. We did not detect such vesicular pRab12 staining in basal ganglia or brainstem cholinergic neurons, and we also did not find a similar staining pattern in non-cholinergic neurons across all brain regions examined. We cannot exclude that the pRab12 staining reflects a phospho-epitope that is distinct from pRab12, and staining of cholinergic forebrain neurons from G2019S-LRRK2-KI mice deficient in Rab12 will be required to address this issue. In addition, the presence of pRab12 on vesicular structures suggests a certain type of stress which is unique to cholinergic forebrain neurons in G2019S-LRRK2-KI mice, and our future work will seek to analyze ciliary deficits and pRab12 accumulation in other mouse models of PD to identify a common nature of endosomal/lysosomal stress which may give insights into mechanisms underlying sporadic PD (Alessi et al., 2024). In contrast to what we observed in basal forebrain cholinergic neurons, the age-dependent loss of cilia in basal ganglia and brainstem cholinergic neurons suggests a different underlying mechanism related to ciliary stability, and our future studies will focus on characterizing the mechanisms underlying those defects.

The different age-dependent onset of the ciliary defects correlates with temporally and anatomically distinct cholinergic innervation deficits. As sampled in the dorsal hippocampus, axons derived from forebrain cholinergic neurons display a dystrophic phenotype which is already present in young G2019S-LRRK2-KI mice. This is consistent with a report showing deficits in cholinergic innervation and executive function already in young mice (Hussein et al., 2022). In human, PET and MRI imaging show a hypercholinergic state in LRRK2-PD patients as well as in asymptomatic LRRK2 mutation carriers. This is unique to the LRRK2 mutation, as not observed in sporadic PD patients or healthy controls, and may reflect a compensatory state to prevent cholinergic deficits mediated by pathogenic LRRK2. Consistent with such compensatory mechanism, LRRK2-PD patients have better olfactory scores and better performance on cognitive tests as compared to sporadic PD patients (Saunders-Pullman et al., 2011; Saunders-Pullman et al., 2014; Somme et al., 2015; Srivatsal et al., 2015), and both olfaction and cognitive features are subject to modulation by central cholinergic integrity (Bohnen et al., 2010). However, such compensatory mechanism may not apply to the G2019S-LRRK2-KI mouse model (Hussein et al., 2022), and in future work it will be important to determine cognitive and olfactory features in G2019S-LRRK2-KI mice and their possible relationship to the early anatomical cholinergic deficits as described here. In contrast to basal forebrain cholinergic neurons, we observed a deficit in projections from brainstem cholinergic neurons in middle-aged but not young animals, and cholinergic cell loss in the PPN in middle-aged animals. Lesioning of ChaT+ neurons in the PPN impairs gait performance in rodents (Chambers et al., 2021), and gait disturbances are common in LRRK2-PD and sporadic PD patients (Chambers et al., 2020; Mirelman et al., 2013). In future studies, it will be important to determine whether middle-aged and aged G2019S-LRRK2-KI mice display deficits in gait performance. In addition, future work is warranted to determine whether ciliary defects in distinct cholinergic brain nuclei are also present in postmortem brains from human LRRK2-PD and sporadic PD patients as compared to healthy controls.

Finally, we find age-dependent cholinergic cell loss in the NBM, PPN and DB in G2019S-LRRK2-KI mice. Strikingly, this is identical to human sporadic PD, where cholinergic cell loss is apparent in the NBM and PPN but not other forebrain or brainstem cholinergic nuclei as analyzed here (Giguère et al., 2018; Surmeier et al., 2017). Few postmortem studies also report cell loss in the DB, even though this region is not part of routine diagnostic sampling protocols (A. K. L. Liu et al., 2018). Since all cholinergic neurons display a ciliary defect with age in the G2019S-LRRK2-KI mice, it will be important to determine whether the particular cholinergic neurons that are vulnerable to cell death require appropriate ciliary signaling for long-term survival.

In summary, pathogenic LRRK2 has a profound effect on cilia in the intact mouse brain which is not limited to cholinergic striatal interneurons, but also observed in all major cholinergic forebrain and brainstem nuclei. Lack of cilia in cholinergic forebrain neurons in mutant LRRK2 mice may be due to a specific type of stress, causing vesicular pRab12 accumulation and followed by the formation of a protein complex which interferes with *de novo* cilia formation as previously described in cells *in vitro*. In contrast, the age-dependent loss of cilia in basal ganglia and brainstem cholinergic neurons may reflect LRRK2-mediated effects on ciliary stability that will require further investigation. Defects in innervation derived from forebrain and brainstem cholinergic neurons show a similar temporal pattern of onset as the ciliary defects and imply wide-ranging defects in cholinergic signaling, with potential impacts on prodromal and early PD phenotypes. The death of a subset of cholinergic neurons in G2019S-LRRK2-KI mice highlights the possibility that ciliary signaling may be important for the long-term survival of those cholinergic neurons that show cellular vulnerability in human PD.

## Methods

### Transgenic mice

Heterozygous G2019S-LRRK2 knock-in (KI) breeding pairs of mice which had been extensively backcrossed to wildtype C57BL/6J mice were obtained from Heather Melrose (Mayo Clinic) (Yue et al., 2015). Heterozygous mice were crossed to yield wildtype and homozygous G2019S-LRRK2-KI progeny. Homozygous G2019S-LRRK2-KI mice were backcrossed to wildtype C57/BL6J mice every sixth generation to prevent genetic drift. Genotyping was performed from tail clips by a PCR-based strategy utilizing primers as described (Yue et al., 2015) (forward 5’CAGGTAGGAGAACAAGTTTAĆ3, reverse 5’GGGAAAGCATTTAGTCTGAĆ3) to yield a 307 bp band in wt, 383 bp band in homozygous, and both bands in heterozygous mice. In humans, the G2019S-LRRK2 mutation is autosomal-dominant, producing similar disease onset and progression in heterozygous and homozygous G2019S mutation carriers (Paisán-Ruiz et al., 2013). Here, and consistent with previous ciliogenesis studies in LRRK2 mouse models, we used homozygous G2019S-LRRK2-KI and wildtype controls (Dhekne et al., 2018; Khan et al., 2021). Age-dependent alterations were examined in young adult (4-5 month old), middle-aged (13-14 months old) and aged (18-24 months old) mice, with exact age recorded in all cases. Both male and female mice were employed for all experiments, with no sex-specific differences observed. Animals were kept in specific pathogen-free conditions at Rutgers New Jersey Medical School and were multiply-housed and maintained on a 12 h light/12 h dark cycle with free access to food and water. All animal procedures were approved by the Rutgers Institutional Animal Care and Use Committee.

### Mouse transcardiac perfusion and brain processing

Transcardiac perfusion was performed with cold PBS for 5 min, followed by cold 4% (v/v) paraformaldehyde (PFA) in PBS for 10 min. Brains were postfixed in 4% PFA overnight at 4°C, immersed in 15% (w/v) sucrose in PBS overnight at 4°C, followed by immersion in 30% (w/v) sucrose in PBS at 4°C overnight or until the brains settled to the bottom of the tube. Brains were then frozen on dry ice, embedded in plastic blocks with OCT (Sakura, #4583), and stored at –80°C. Embedded brains were oriented to cut coronal sections and serially sectioned at –20°C using a Leica CM3050 S cryostat. Sections (30 μm) were collected and submerged in PBS, and free-floating sections kept at 4°C and transferred into fresh PBS 24 h later. All stainings were done with fresh cryosections whenever possible, and all remaining sections stored in cryoprotectant solution (30% (w/v) sucrose, 30% (v/v) ethylene glycol, 0.1% (w/v) sodium azide in PBS) at –20°C.

### Immunofluorescence staining

Mouse brain primary cilia detection was performed as described (Dhekne et al., 2018; Khan et al., 2024, 2021) with the following modifications. Free-floating coronal 30 μm sections were transferred to 24-well plates and quenched in 0.3% (w/v) Sudan Black (Sigma Aldrich, #199664) in 70% ethanol for 10 min, followed by washing 3 x 10 min in PBS at RT. Sections were permeabilized and blocked in 5% normal donkey serum (NDS) (Sigma Aldrich, #D9663) in 0.3% Triton X-100/PBS for 2 h at RT. For each brain region of interest, two non-consecutive sections were co-stained with primary antibodies against adenylate cyclase III (ACIII) and choline acetyltransferase (ChaT) for 48 h in 5% NDS in 0.3% Triton X-100/PBS at 4°C. Sections were washed 3 x 10 min with 0.3% Triton X-100/PBS and incubated with the respective highly cross-absorbed Alexa Fluor secondary antibodies in 5% NDS in 0.3% Triton X-100/PBS for 2 h at RT. Slices were then washed for 4 x 10 min with 0.3% Triton X-100/PBS and incubated in a DAPI containing dye solution (NucBlueTM Fixed Cell Stain ReadyProbes TM reagent, Invitrogen, #R37606; 1 drop/ml in PBS) for 30 min, followed by mounting on glass slides, overlayed with a hardening mountant and coverslip, and stored at 4°C in the dark. Cilia staining in striatal fast-spiking interneurons was performed as described above with anti-ACIII and either anti-parvalbumin or anti-calretinin antibodies, respectively. All antibodies and reagents employed are listed in **Table 1**, and Bregma coordinates for all brain regions sampled are listed in **Table 2**.

**Table 2.**
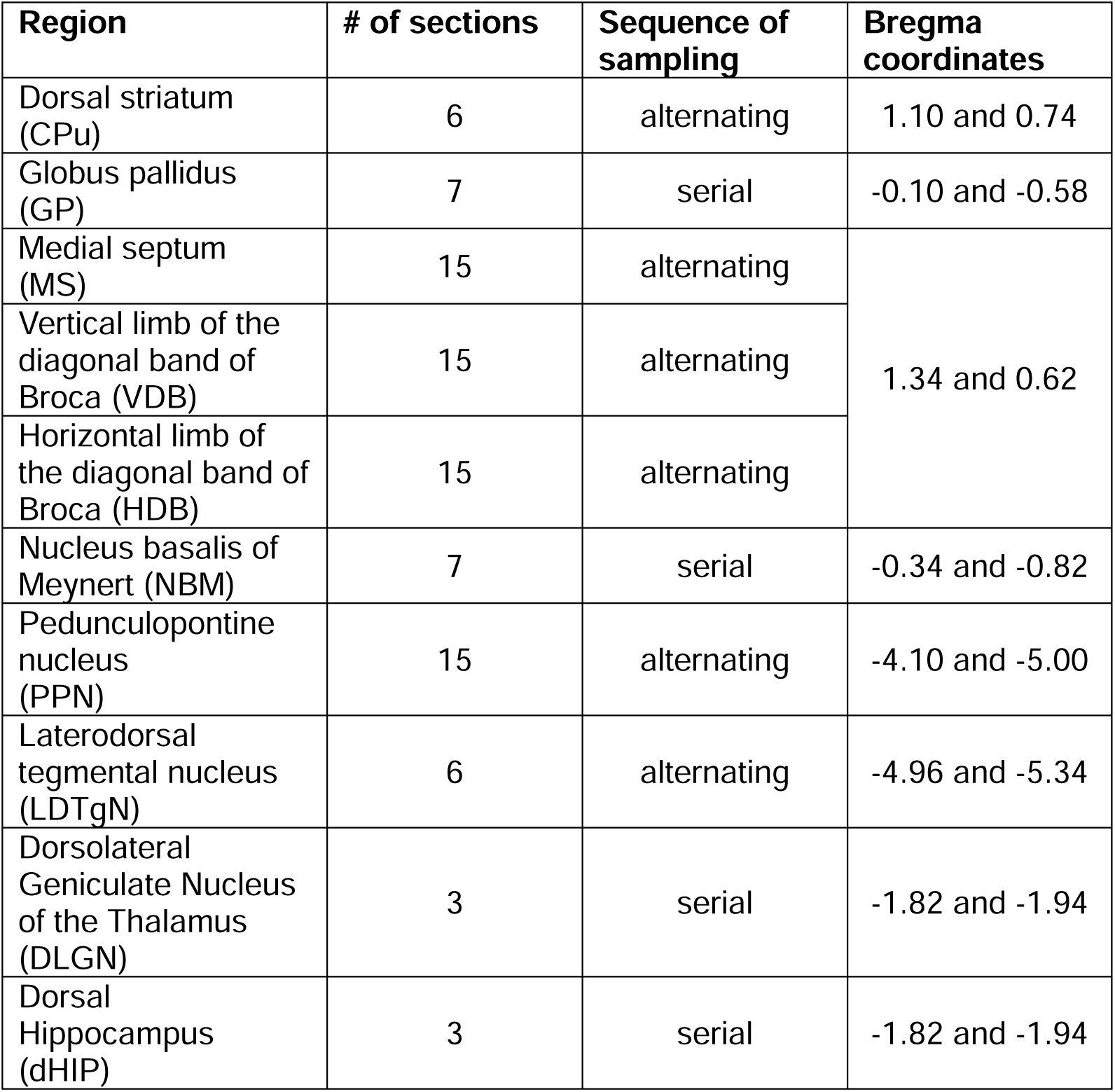
Regional sampling parameters.

Phospho-Rab12 and ChaT co-staining was performed as described above. To assess the phospho-state-specific nature of the staining, sections were quenched in 0.3% (w/v) Sudan Black in 70% ethanol for 10 min at RT, followed by washing 3 x 10 min in PBS at RT. Sections were incubated for 10 min at RT in 1xNEBuffer with 1 mM MnCl_2_ provided by the Lambda Protein Phosphatase kit, followed by incubation in 1x NEBuffer with 1mM MnCl_2_ for 6 h at 37 °C in either the presence or absence of 10,000 units/ml Lambda phosphatase. Sections were then washed in PBS, followed by blocking for 2 h at RT and staining as described above.

For better visualization of pRab12-positive vesicular structures, an antigen retrieval protocol was employed. Sections were washed 2 x 5 min in PBS, blocked with 5% NDS in 0.3% Triton X-100/PBS for 30 min at RT, followed by 2 x 5 min washes in PBS and permeabilization with 0.5% Triton X-100/PBS for 15 min at RT. Before antigen retrieval, sections were washed 2 x 5 min in PBS. Tris-based antigen unmasking solution (Vector Laboratories, #H-3301) was diluted 1:1000 in ddH_2_O and microwaved 10-20 sec until boiling, and sections incubated in this solution at 100 °C on a dry bath for 30 min. Sections were allowed to cool down to RT, and then washed 2 x 5 min with PBS at RT before blocking with 5% NDS in 0.3% Triton X-100/PBS for 24 h at 4°C. Slices were washed 2 x 5 min with PBS at RT and incubated with primary antibodies against pRab12 and ChAT in 1% NDS in 0.1% Triton X-100/PBS for 48 h at 4°C. After primary antibody incubation, sections were washed 2 x 5 min with PBS and incubated with the respective highly cross-absorbed Alexa Fluor secondary antibodies in 1% NDS in 0.1% Triton X-100/PBS for 1 h at RT. Sections were washed 3 x 10 min with PBS, incubated in a DAPI nuclear dye solution and mounted as described above.

For determination of cell counts, sections were stained as described above. Sections encompassing globus pallidus were co-stained with antibodies against ChaT and calbindin (Geula et al., 1993) for more precise definition of this brain area. Basal forebrain sections were co-stained with anti-ChaT and anti-p75, and brainstem sections with anti-ChaT and anti-nNOS, respectively (Boskovic et al., 2019; Geula et al., 1993).

Vesicular acetylcholine transferase (VAChT) staining was performed essentially as described (Hussein et al., 2022). Three consecutive coronal sections which include both dorsal hippocampus and DLGN were quenched in Sudan black, washed in PBS, permeabilized with 0.25% Triton X-100/PBS for 15 min, and blocked in 10% bovine serum albumin (BSA) (GOLDBIO, #A-421-100) in PBS for 1 h at RT. Sections were incubated with primary antibody against VAChT in 2% BSA in 0.1% Triton X-100/PBS for 72 h at 4°C. Sections were washed 3 x 10 min with PBS and incubated with highly cross-absorbed Alexa Fluor secondary antibody in 2% BSA in 0.1% Triton X-100/PBS for 1 h at RT. Sections were washed 3 x 10 min with PBS, incubated in a DAPI nuclear dye solution and mounted as described above.

VAChT and ChAT co-staining was performed as follows. Slices were quenched in Sudan black, washed in PBS, permeabilized with 0.25% Triton X-100/PBS for 15 min, and blocked in 5% BSA and 5% NDS in PBS for 1 h at RT. Sections were incubated with primary antibodies against VAChT and ChAT in 2% BSA and 3% NDS in 0.1% Triton X-100/PBS for 72 h at 4°C. Sections were washed 3 x 10 min with PBS and incubated with highly cross-absorbed Alexa Fluor secondary antibodies in 2% BSA and 3% NDS in 0.1% Triton X-100/PBS for 1 h at RT. Sections were washed 3 x 10 min with PBS, incubated in a DAPI nuclear dye solution and mounted as described above.

### Image acquisition

All images were acquired on an Olympus FV1000 Fluoview microscope using a 60 x 1.2 NA water objective (UPlanSApo) for ciliogenesis, pRab12 staining and VAChT staining in DLGN, a 20 x 0.4 NA objective to determine dystrophic neuronal projections in the dorsal hippocamal CA1/CA2 region, and a 10 x 0.30 NA objective for determination of cell counts. All images were collected using single excitation for each wavelength separately and dependent on secondary antibodies, and were acquired with the same laser intensities, acquisition settings and with a step size of 1 μm.

For cilia determinations, two non-consecutive sections were analyzed for each brain region (**Table 2**). Given the region-specific differences in ChaT+ cell density, determination of ciliogenesis and quantification of pRab12 staining was performed over 15-40 independent images per section.

For determining innervation density in the DLGN, 5 random areas per section from 3 consecutive sections were imaged, with single image dimensions of 212 μm x 212 μm. For detecting dystrophic axons, 3 adjacent non-overlapping images were acquired in CA1/CA2 region of dorsal hippocampus, with single image dimensions of 636 μm x 636 μm.

Image acquisition for determination of ChaT+ cell counts across different brain areas was performed as described (Pappas et al., 2018, 2015) with minor modifications. For dorsal striatum (caudate putamen), 6 alternating serial 30 μm sections were imaged. For cell counts in globus pallidus and NBM, images from 7 serial 30 μm sections were acquired. For medial septum, vertical and horizontal diagonal bands, 15 alternating serial sections starting with the beginning of the medial septum as determined by the presence of ChaT+ neurons were acquired. The corpus callosum and lateral ventricle were used as anatomical identifying boundaries, and sections compared to mouse brain atlas to define approximate boundaries between medial septum and vertical diagonal band. For the pedunculopontine nucleus, 15 alternating serial sections starting with the beginning of the nucleus as determined by ChaT+ counts were imaged. Given previous reports of sparse distribution of ChaT neurons in the rostral as compared to the caudal rat PPN (Martinez-Gonzalez et al., 2011; Mena-Segovia et al., 2009), cell counts were also separately determined for rostral (first 6 alternating sections) and caudal (last 6 alternating sections) PPN. For laterodorsal tegmental nucleus, 6 alternate serial 30 μm sections were acquired. Number of sections acquired for each brain region and Bregma coordinates are indicated in **Table 2**.

### Image analysis

Image analysis was performed using the Fiji by ImageJ software, the MATLAB hosted ACDC software (Lauring, Cilia, 2019), or Imaris v 10.0.1 software. To analyze primary cilia, images were individually analyzed across all z-stacks by an observer blind to genotype and age, and cells scored as ciliated when the cilium was encompassed within ChaT+ cell bodies. For cilia determination from ChaT-cells, cells were scored as ciliated when nuclear DAPI stain overlapped with a cilium in the same z-stack, and percent ciliation determined by counting the number of ciliated nuclei over total nuclei.

Cilia length analysis was performed using ACDC software (Lauring et al., 2019). 20-30 ChaT+ cells from the dorsomedial CPu were scored per section over two independent sections per animal. Cilia counts based on ACDC software were comparable to those obtained by the Fiji-based approach.

For cilia determination in ChaT+ neurons in intermediate CPu, the area was subdivided into four equal quadrants, and ciliogenesis quantified separately from dorsomedial and ventrolateral areas, respectively. Ciliogenesis was determined from ChaT+ neurons in both MS and VDB. In some cases, ciliogenesis was additionally quantified from ChaT+ neurons in HDB, with no differences in ciliogenesis observed between these three areas. Similarly, in some cases, ciliogenesis was scored from ChaT+ neurons in rostral versus caudal PPN, with no differences observed between these two areas. For each section and brain area, 50-100 ChaT+ neurons and 200-400 ChaT-neurons were scored, and average ciliation over two sections yielded one datapoint per mouse per brain area (Dhekne et al., 2018; Khan et al., 2021).

The number of pRab12+ structures per ChaT+ cell was manually determined in Fiji from 50-100 ChAT+ cells over 2 independent sections per animal. Quantification of pRab12 intensity was performed in Fiji over non-saturated images acquired with the same laser intensities and acquisition settings. We first generated a region of interest (ROI) by drawing a contour around a ChAT+ cell. Using the drawn contour, we moved it to an adjacent area with no pRab12 signal and measured the mean gray value. Using this, we realigned the ROI with the ChAT+ cell and background subtracted the mean value to account for differences in tissue background staining between animals. We then used the thresholding function and measured the integrated density for each cell. This value is the product of the area of the ROI and mean gray value. Since this value is driven by the area of each ROI, we confirmed that there were no significant differences in cell size as measured by the mean ROI area between our experimental groups. 20-45 ChAT+ cells were scored over 2 slices per animal.

VAChT+ dystrophic axons in dorsal hippocampal CA1/CA2 regions were quantified from maximum intensity projections of z-stacks from three 20x images per section, three sections per animal and 5 animals per age and genotype. We first set a threshold and generated a mask that was used to analyze the suprathreshold particles within the ROI. A mask was used to quantify particles larger than 3 μm^2^. The threshold and mask remained constant for all images quantified in a section. Using the Analyze Particles function, we measured cluster number over 1908 μm x 1908 μm area in the CA1 and CA2 regions of the dorsal hippocampus from 5 animals per genotype and age.

VAChT+ axonal density in DLGN was determined from maximum intensity projections of z-stacks from 5 60x images per section, 3 sections per animal, 5 animals per age and genotype as previously described (Hussein et al., 2022). Each image was first background subtracted by the average gray value of a nearby area lacking specific VAChT signal. Images were then thresholded using the same threshold settings to generate a mask of the immunofluorescent signal, and the output image subjected to Analyze > Analyze Particles function. For each image, the density of VAChT immunolabeled fibers was expressed as the percent area occupied by suprathreshold pixels.

Automated identification and quantification of ChAT+ cells in distinct brain regions was conducted using the spot detection function in the Imaris v.10.0.1 software as described (Chen et al., 2020; González-Rodríguez et al., 2021). Quantification of ChAT+/p75+ cells in MS, VDB and HDB, and ChAT+/nNOS+ cells in the PPN and LDTgN was conducted using the spot detection function in conjunction with the colocalization function in Imaris software. The number of 30 μm coronal sections encompassing the different forebrain and brainstem nuclei was unchanged across genotypes, indicating that the rostrocaudal dimensions of those nuclei were not altered (Yeo et al., 1997). Hence, alterations in cell counts reflect alterations in total ChaT+ neuronal cell numbers.

## Statistics

Data were plotted using GraphPad Prism 9 software (GraphPad Prism, RRID:SCR_002798). In all cases, error bars indicate SEM, and two-tailed unpaired Student’s T-test was employed to test significance.

## Data availability

All primary data associated with each figure has been deposited in a repository and can be found at https://doi.org/10.5281/zenodo.12601699 and https://doi.org/10.5281/zenodo.12608526.

## Acknowledgements

This work was supported by a Busch Biomedical Research Grant from Rutgers University (to S.H.). We are grateful to Vinita Chitoor (UCSF) for guidance on phosphatase treatment of brain slices, and to Emily Rocha (University of Pittsburgh) for guidance on staining protocols involving endolysosomal markers.

## Figure Legends

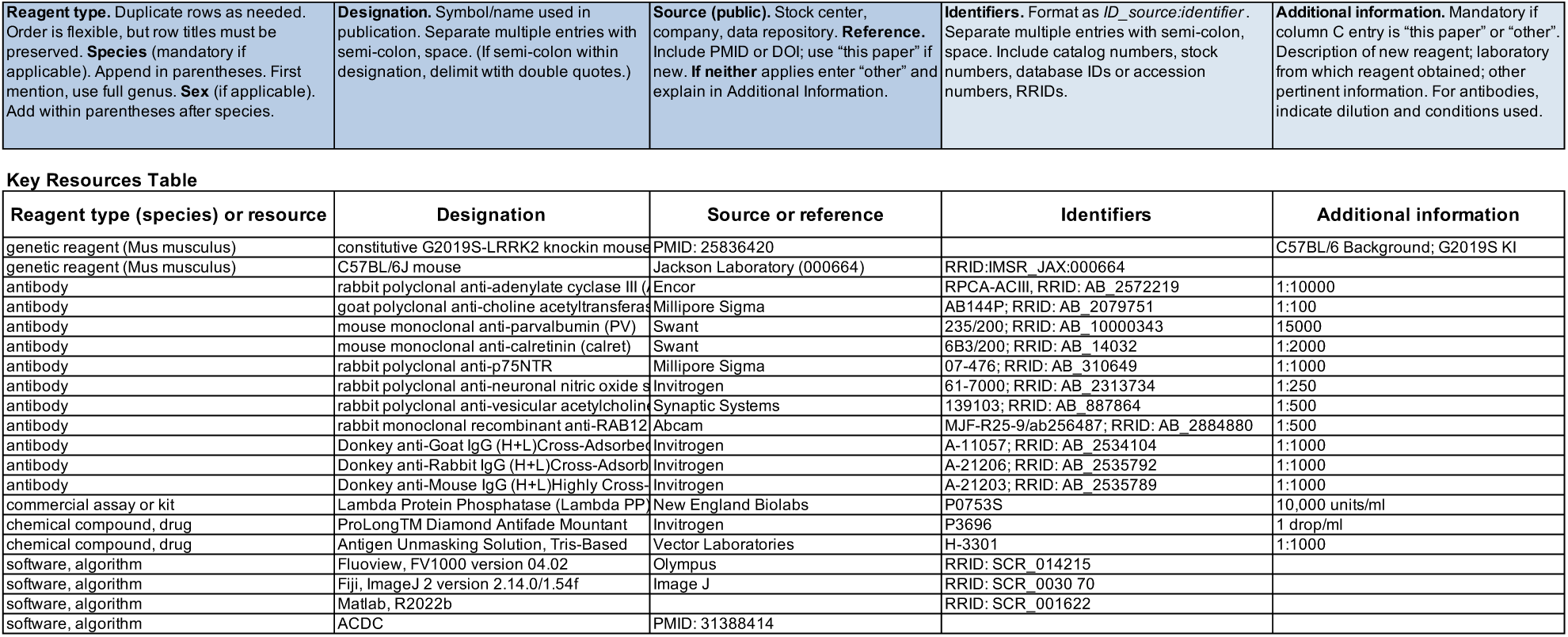

**Figure 1 – Figure supplement 1.**
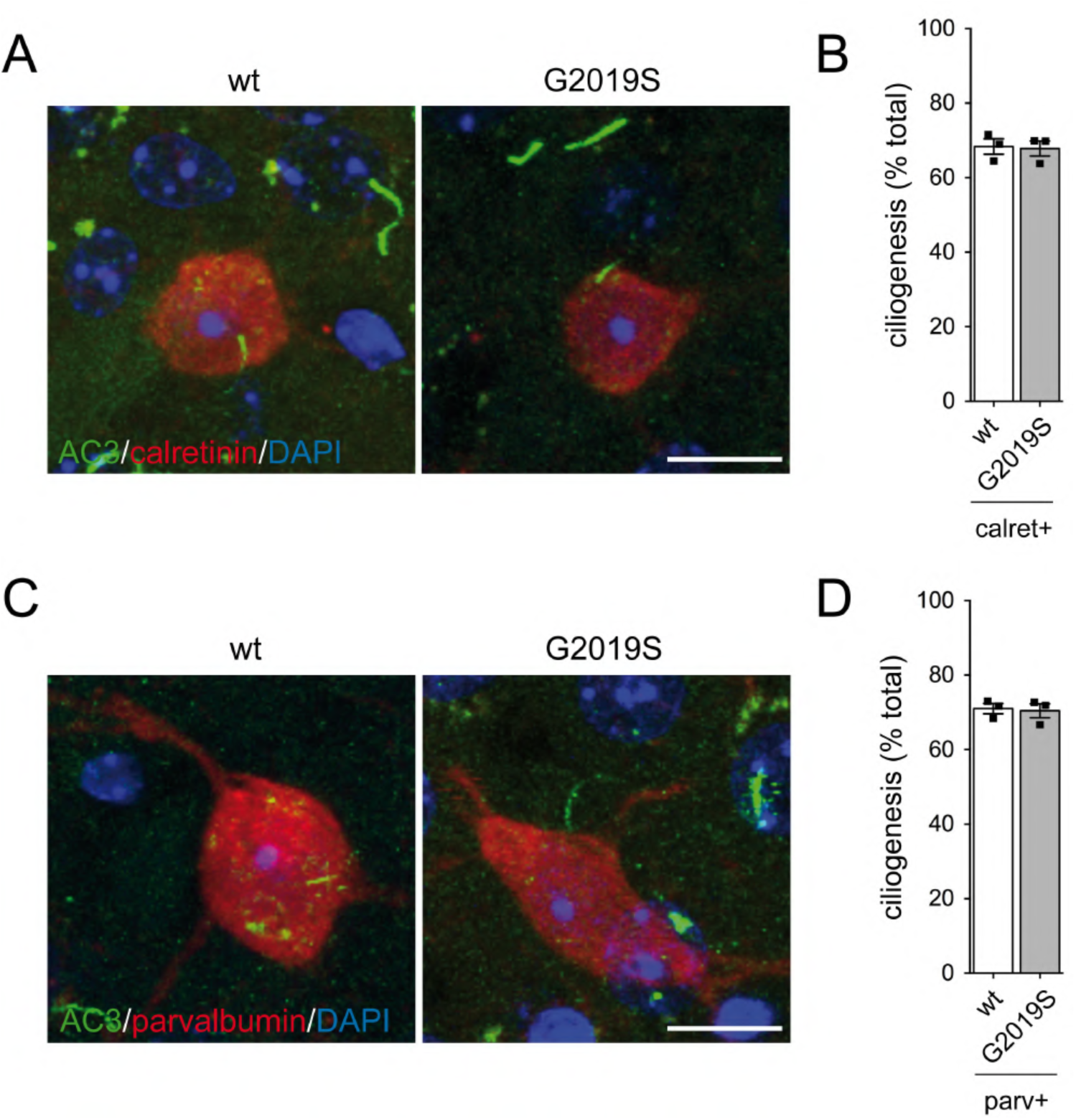
Normal ciliation in calretinin-positive and parvalbumin-positive interneurons in CPu of aged G2019S-LRRK2-KI mice. (**A**) Confocal images of calretinin-positive interneurons in CPu from aged wt and G2019S mice stained for AC3, calretinin and DAPI. Scale bar, 10 μm. (**B**) Quantitation of ciliation in calretinin-positive (calret+) interneurons in CPu from aged (18-24 months) wt and G2019S mice. (**C**) Confocal images of parvalbumin-positive interneurons in CPu from aged wt and G2019S mice stained for AC3, parvalbumin and DAPI. Scale bar, 10 μm. (**D**) Quantitation of ciliation in parvalbumin-positive (parv+) interneurons in CPu from aged (18-24 months) wt and G2019S mice. Each datapoint is from an individual mouse, analyzing 2 sections per animal and 100-120 calret+ or parv+ cells, respectively. Bars represent mean ± s.e.m (n=3).

**Figure 5 – Figure supplement 1.**
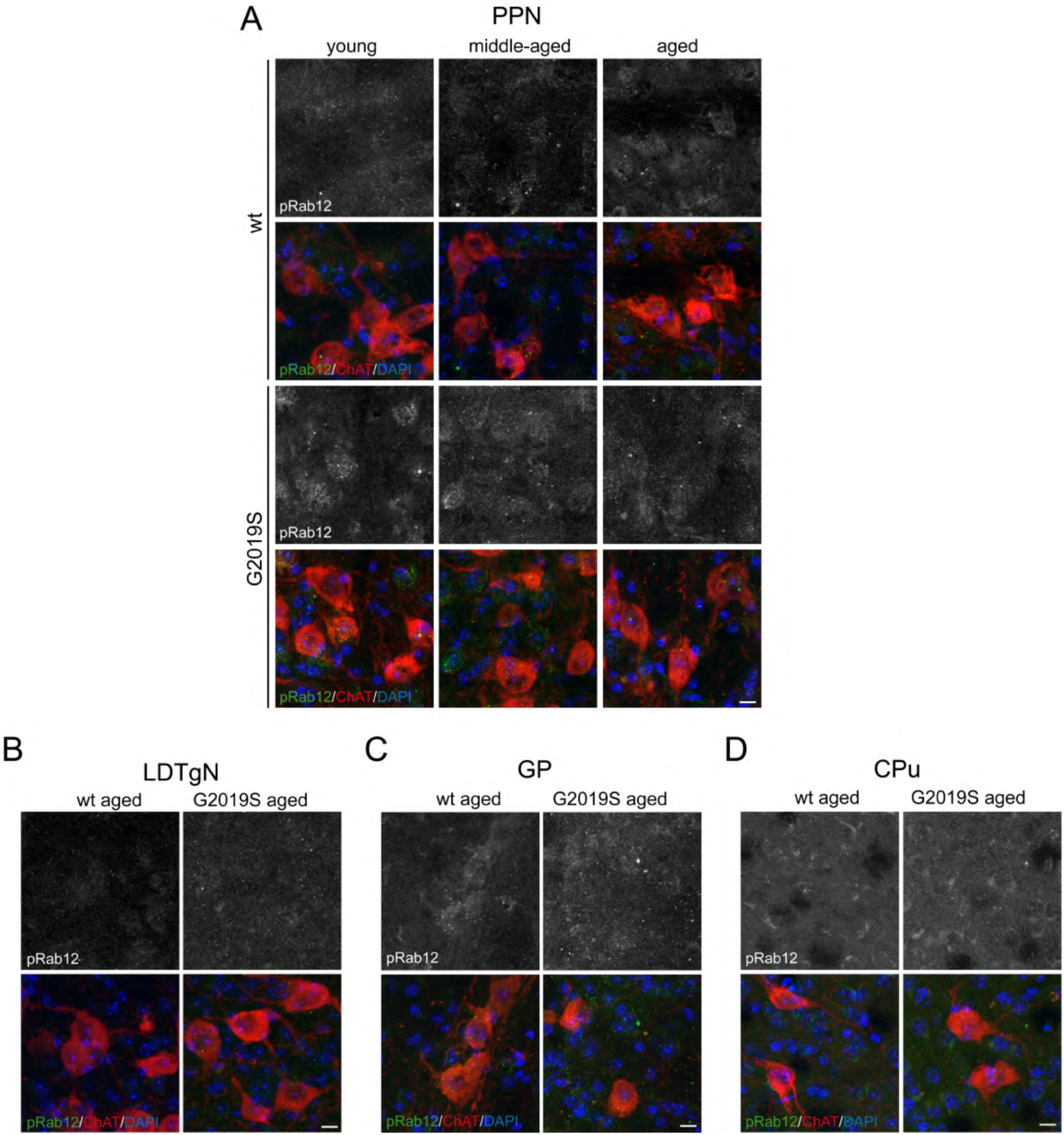
Cholinergic neurons in brainstem and basal ganglia from G2019S-LRRK2-KI mice do not show staining with pRab12 antibody. (**A**) Confocal images of cholinergic neurons in pedunculopontine nucleus (PPN) from young, middle-aged and aged wt and G2019S mice stained with antibodies against phospho-Rab12 (pRab12), choline acetyltransferase (ChaT) and DAPI. (**B**) Cholinergic neurons in LDTgN from aged wt and G2019S mice stained as above. (**C**) Cholinergic neurons in GP from aged wt and G2019S mice stained as above. (**D**) Cholinergic neurons in CPu from aged wt and G2019S mice stained as above. All scale bars are 10 μm.

**Figure 5 – Figure supplement 2.**
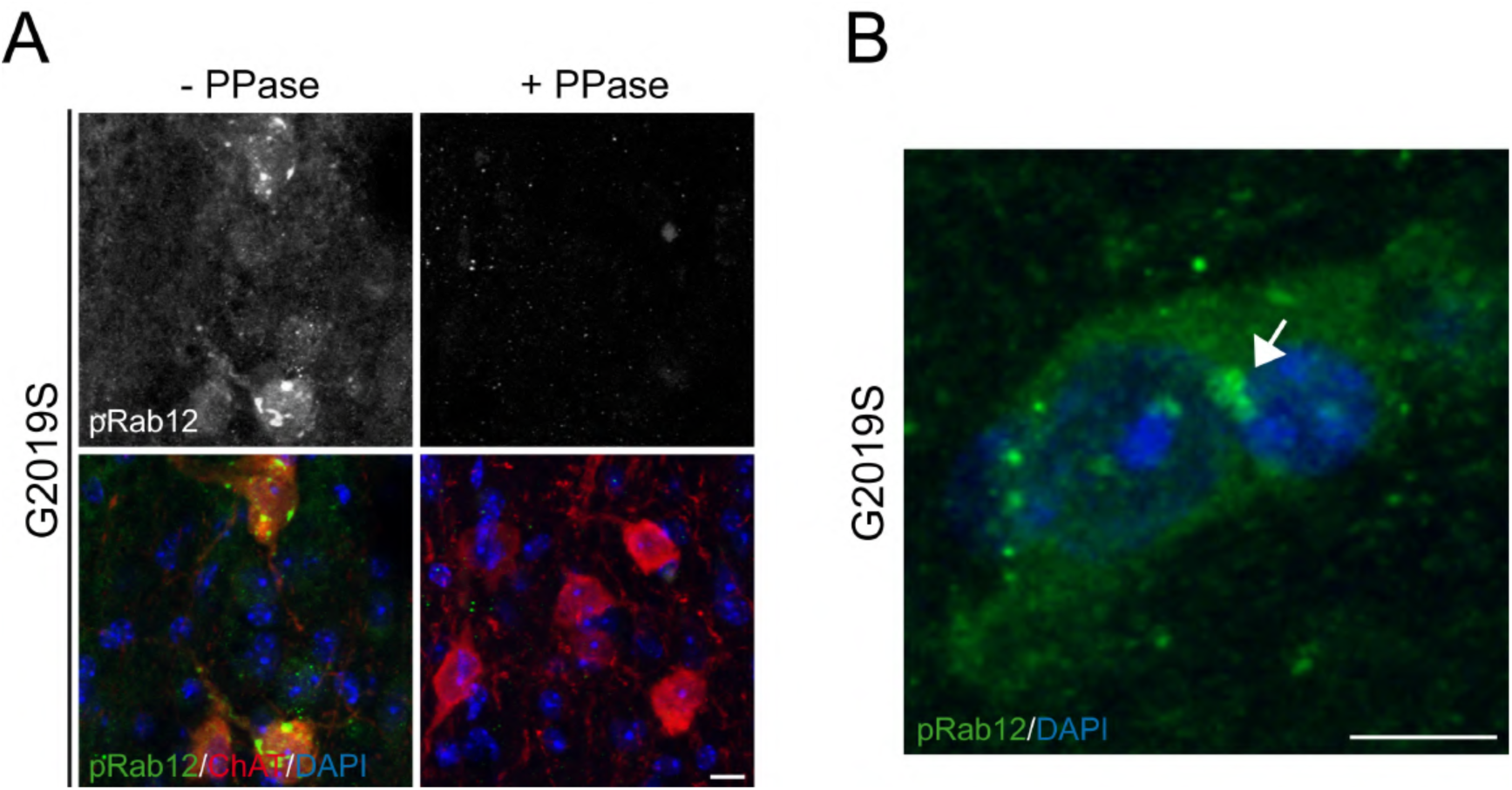
pRab12 staining is phospho-state-specific and vesicular in nature. (**A**) Confocal images of cholinergic neurons in VDB from middle-aged G2019S mice either with or without pretreatment with phosphatase (PPase) before staining with antibodies against phospho-Rab12 (pRab12), choline acetyltransferase (ChaT) and DAPI. Scale bar, 10 μm. (**B**) Confocal image of cholinergic neuron in HDB from middle-aged G2019S mouse stained with pRab12 antibody and DAPI after an antigen retrieval protocol. Arrow points to pRab12-positive vesicular structures. Scale bar, 10 μm.

**Figure 6 – Figure supplement 1.**
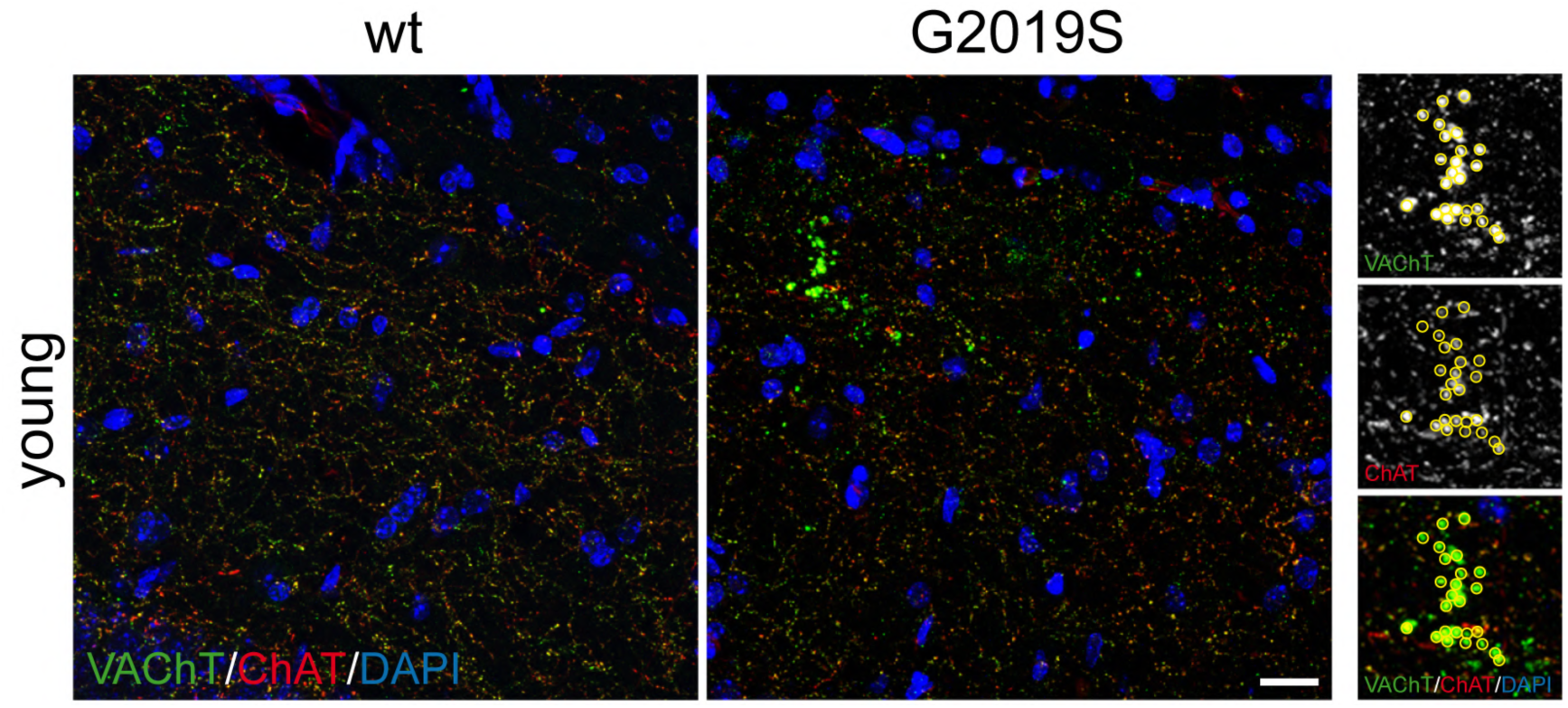
Dystrophic axons derived from basal forebrain cholinergic neurons in G2019S-LRRK2-KI mice stain positive for both VAChT and ChaT. Confocal images of hippocampal CA1/2 regions from young wt and G2019S mice stained for VAChT, ChaT and DAPI. Insert shows colocalization of both cholinergic markers. Scale bar, 100 μm.

**Figure 7 – Figure supplement 1.**
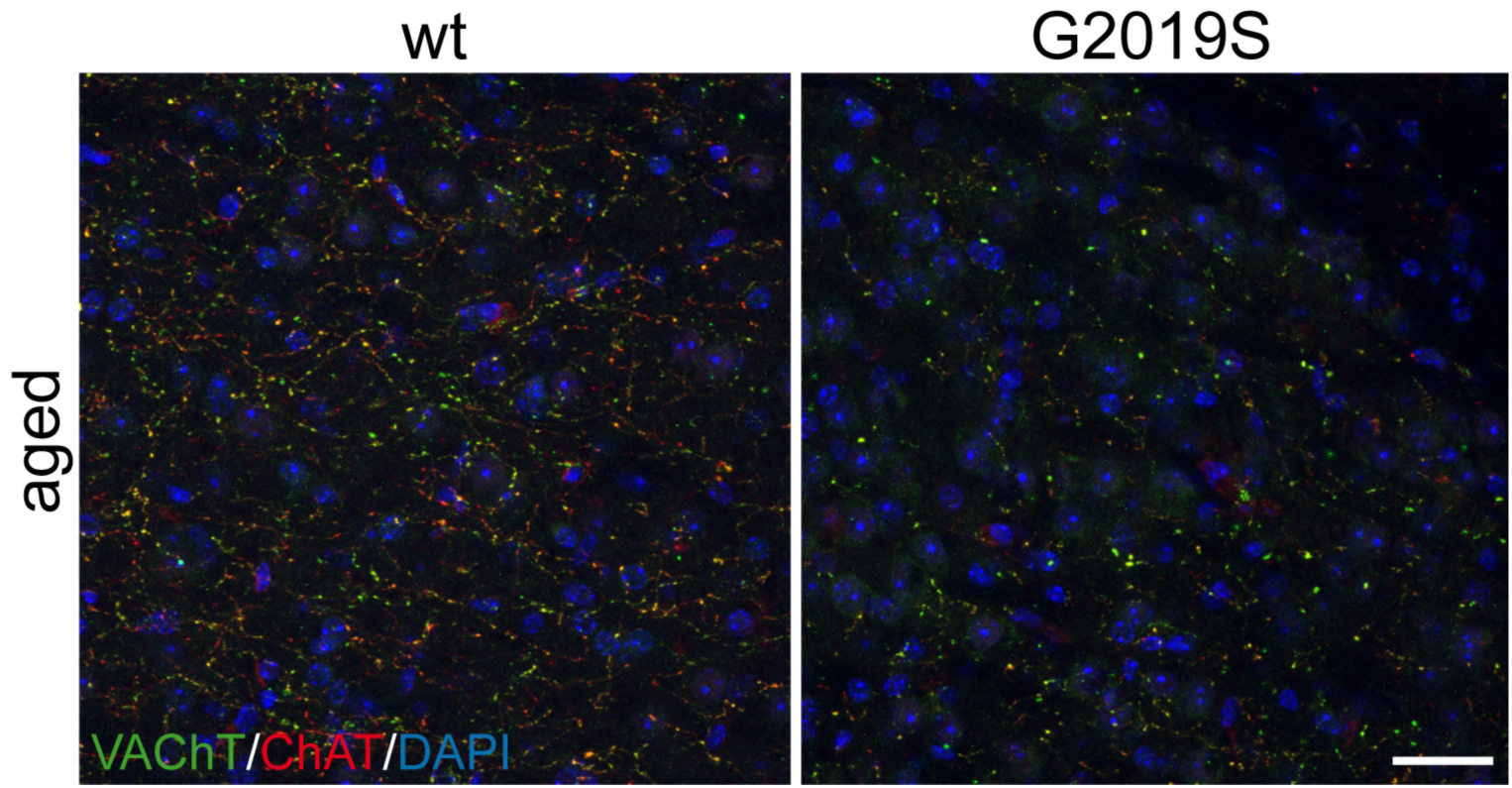
Cholinergic innervation in dLGN from G2019S-LRRK2-KI mice as assessed by both VAChT and ChaT staining. Confocal images of dLGN from aged wt and G2019S mice stained for VAChT, ChaT and DAPI. Scale bar, 30 μm.

**Figure 9 – Figure supplement 1.**
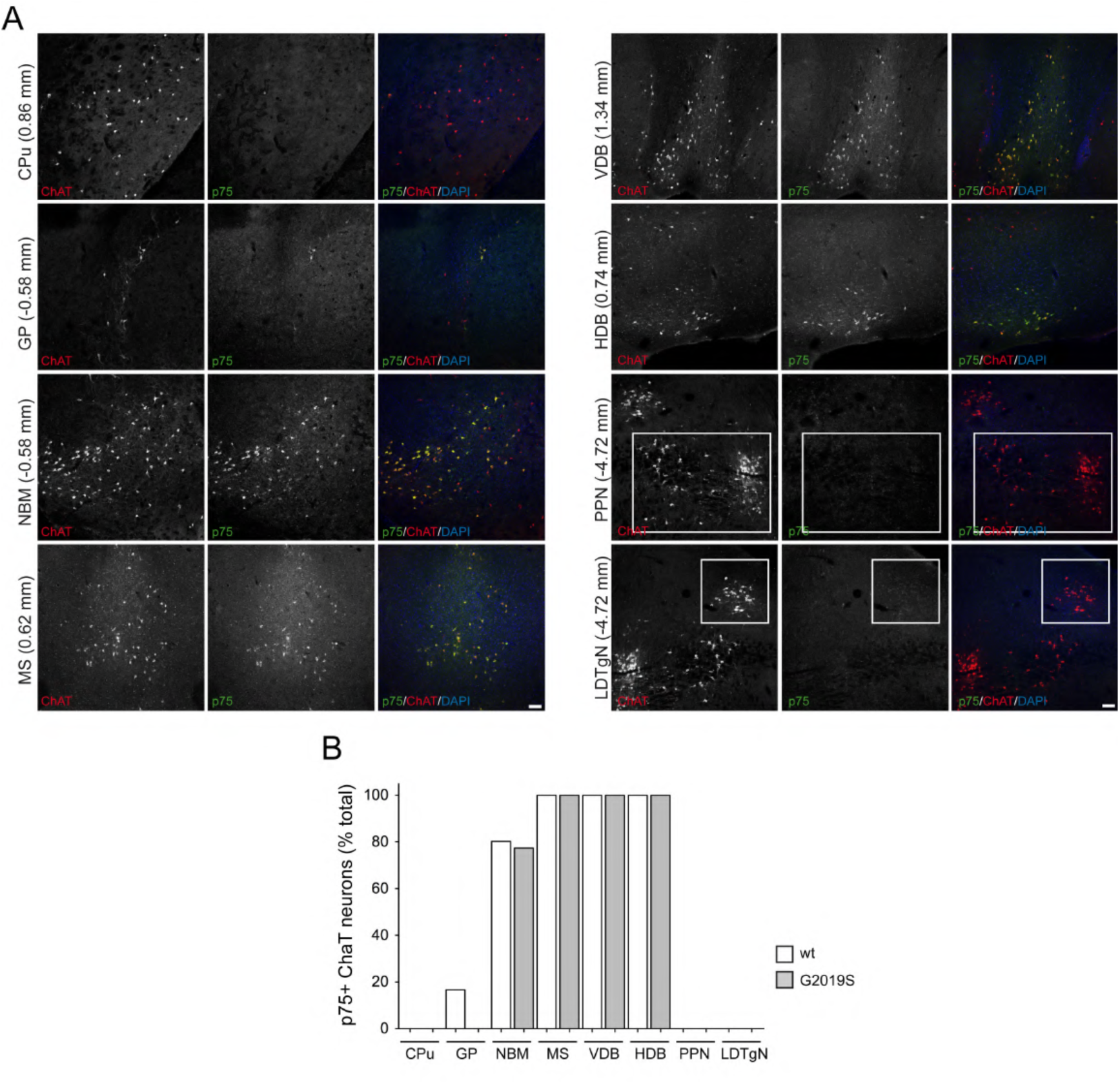
Cholinergic neurons in medial septum and diagonal band are positive for p75. (**A**) Confocal images of different brain areas from 13-months old wt mice stained for ChaT, p75 and DAPI, with Bregma coordinates indicated to the left. Scale bar, 100 μm. (**B**) Quantification of ChaT+ neurons co-stained with p75 from different brain aras as indicated in middle-aged wt and G2019S mice. Colocalization was scored from 50-100 cholinergic neurons per brain area and genotype.

**Figure 9 – Figure supplement 2.**
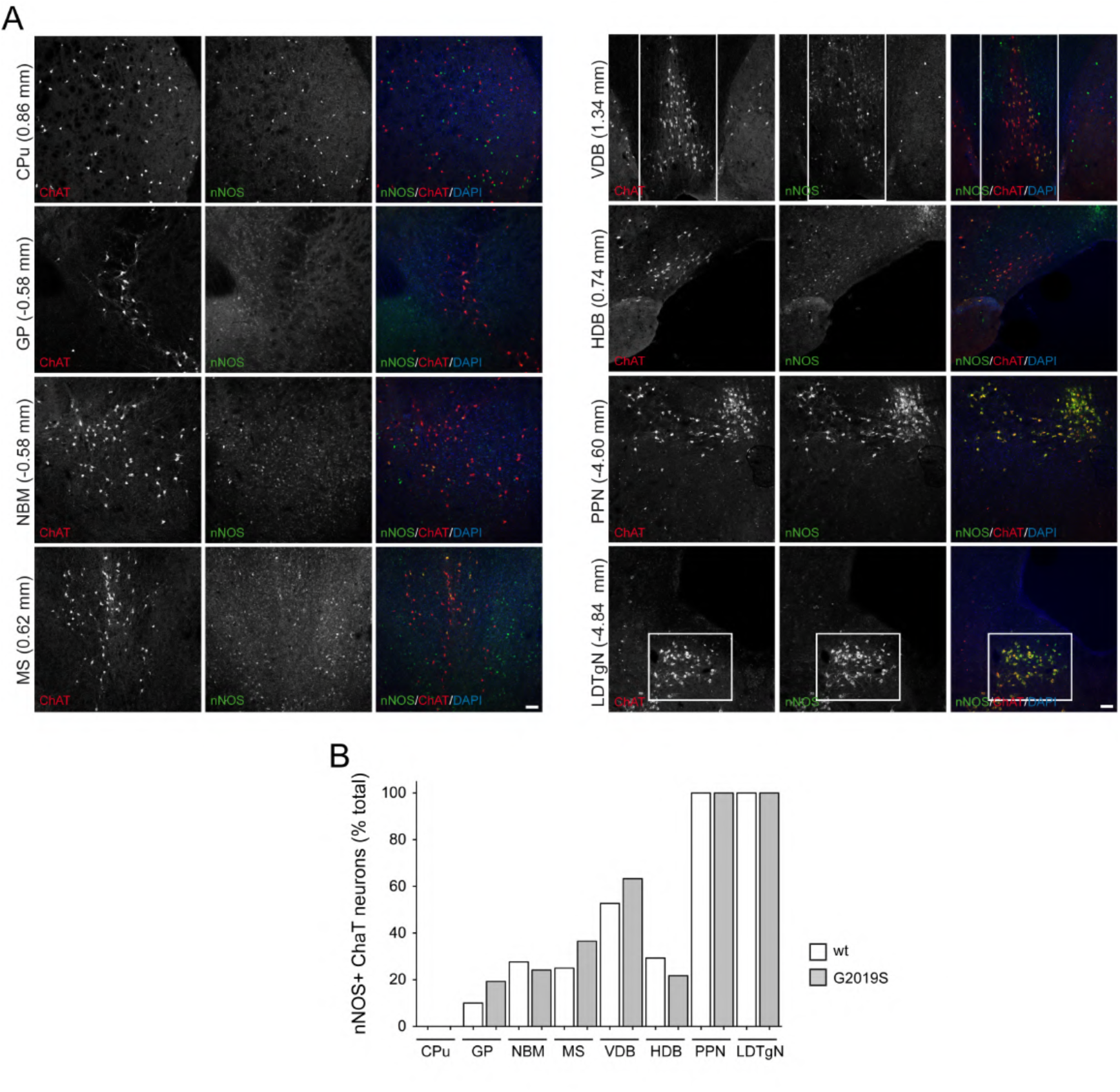
Brainstem cholinergic neurons in PPN and LDTgN are positive for nNOS. (**A**) Confocal images of different brain areas from 13-months old wt mice stained for ChaT, nNOS and DAPI, with Bregma coordinates indicated to the left. Scale bar, 100 μm. (**B**) Quantification of ChaT+ neurons co-stained with nNOS from different brain aras as indicated in middle-aged wt and G2019S mice. Colocalization was scored from 50-100 cholinergic neurons per brain area and genotype.

**Figure 9 – Figure supplement 3.**
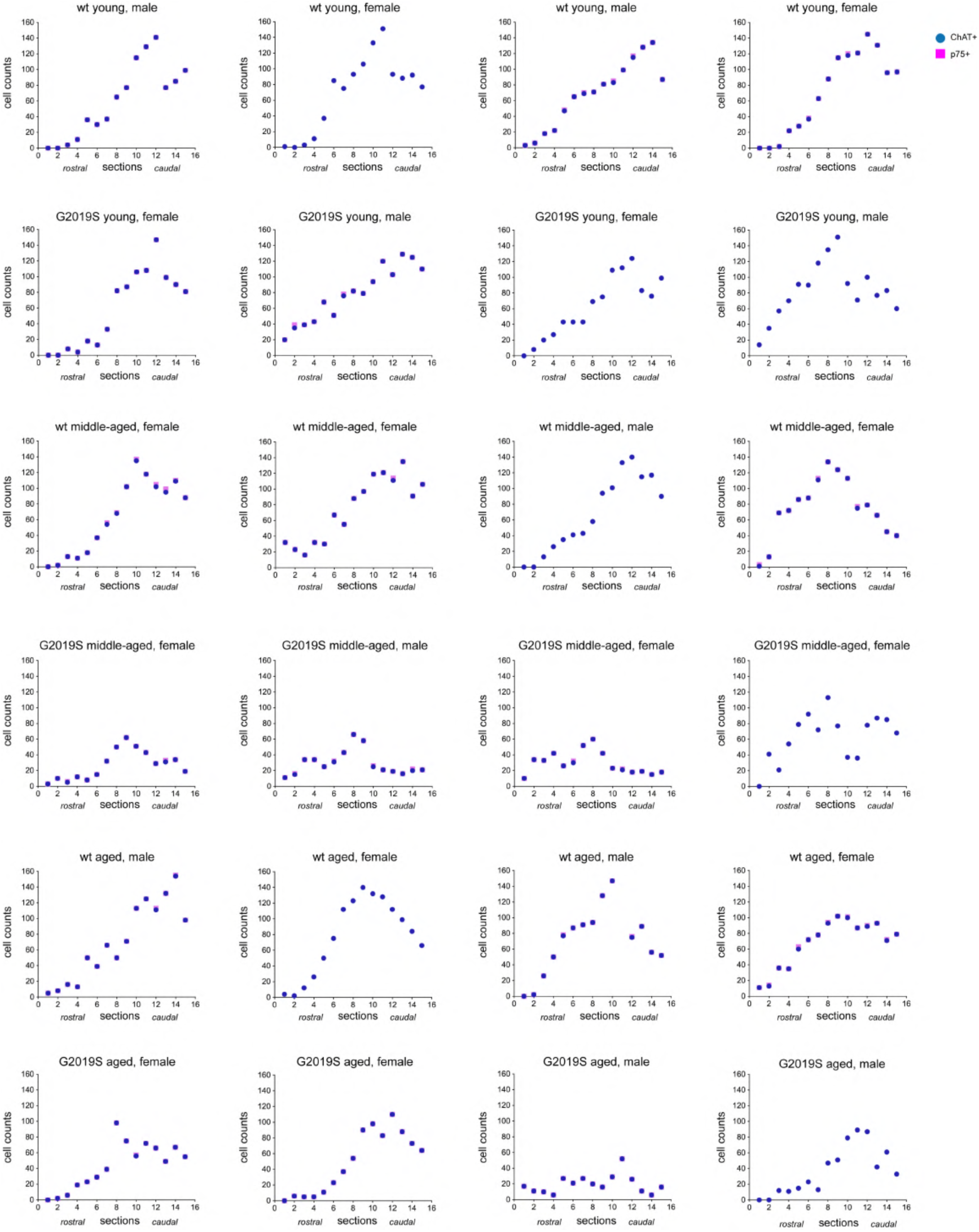
Cholinergic cell counts in HDB. ChaT+ and p75+ cell counts in alternate sections through the horizontal diagonal band from young adult (4-5 months), middle-aged (13-14 months) and aged (18-24 months) male and female mice as indicated.

**Figure 9 – Figure supplement 4.**
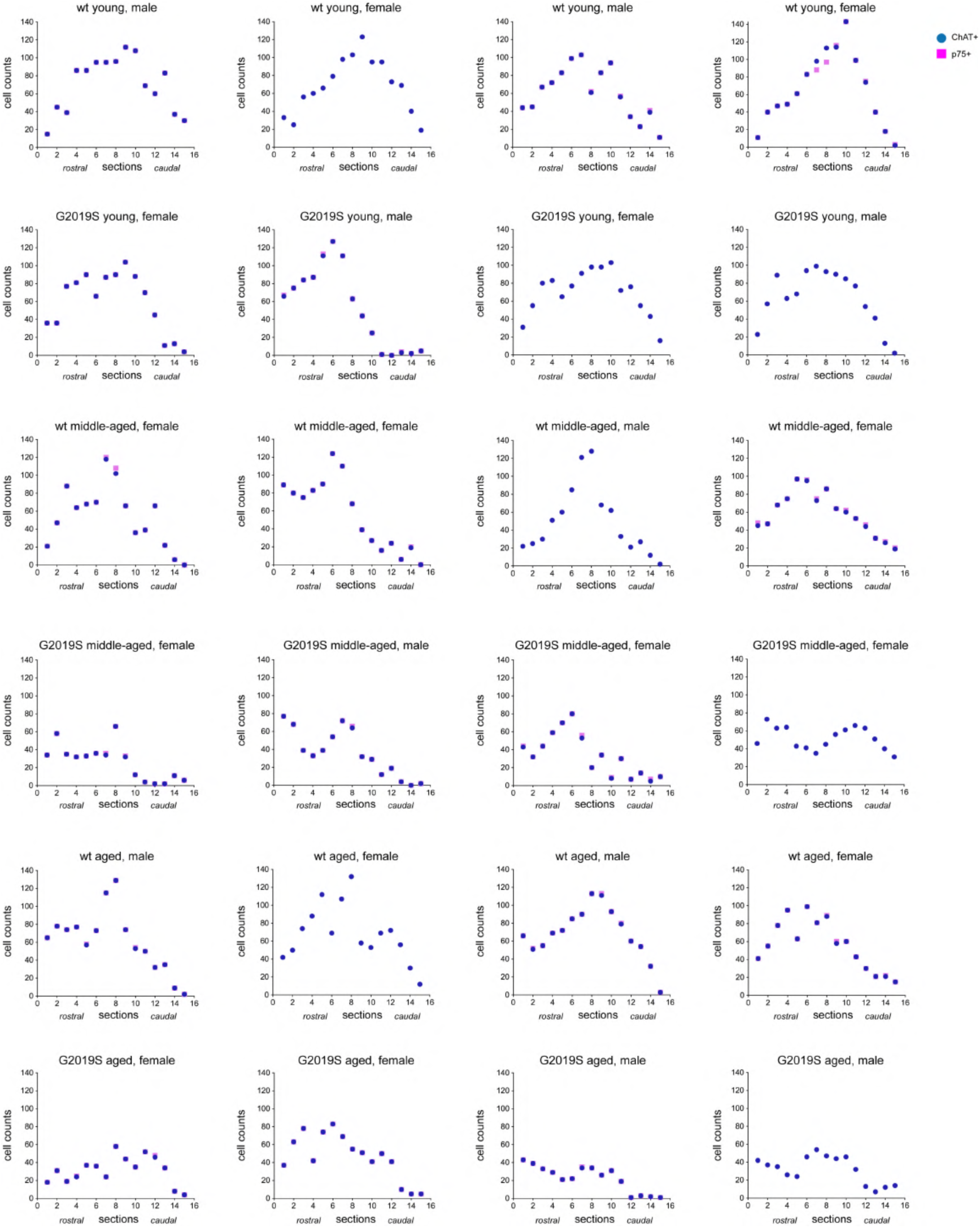
Cholinergic cell counts in VDB. ChaT+ and p75+ cell counts in alternate sections through the vertical diagonal band from young adult (4-5 months), middle-aged (13-14 months) and aged (18-24 months) male and female mice as indicated.

**Figure 9 – Figure supplement 5.**
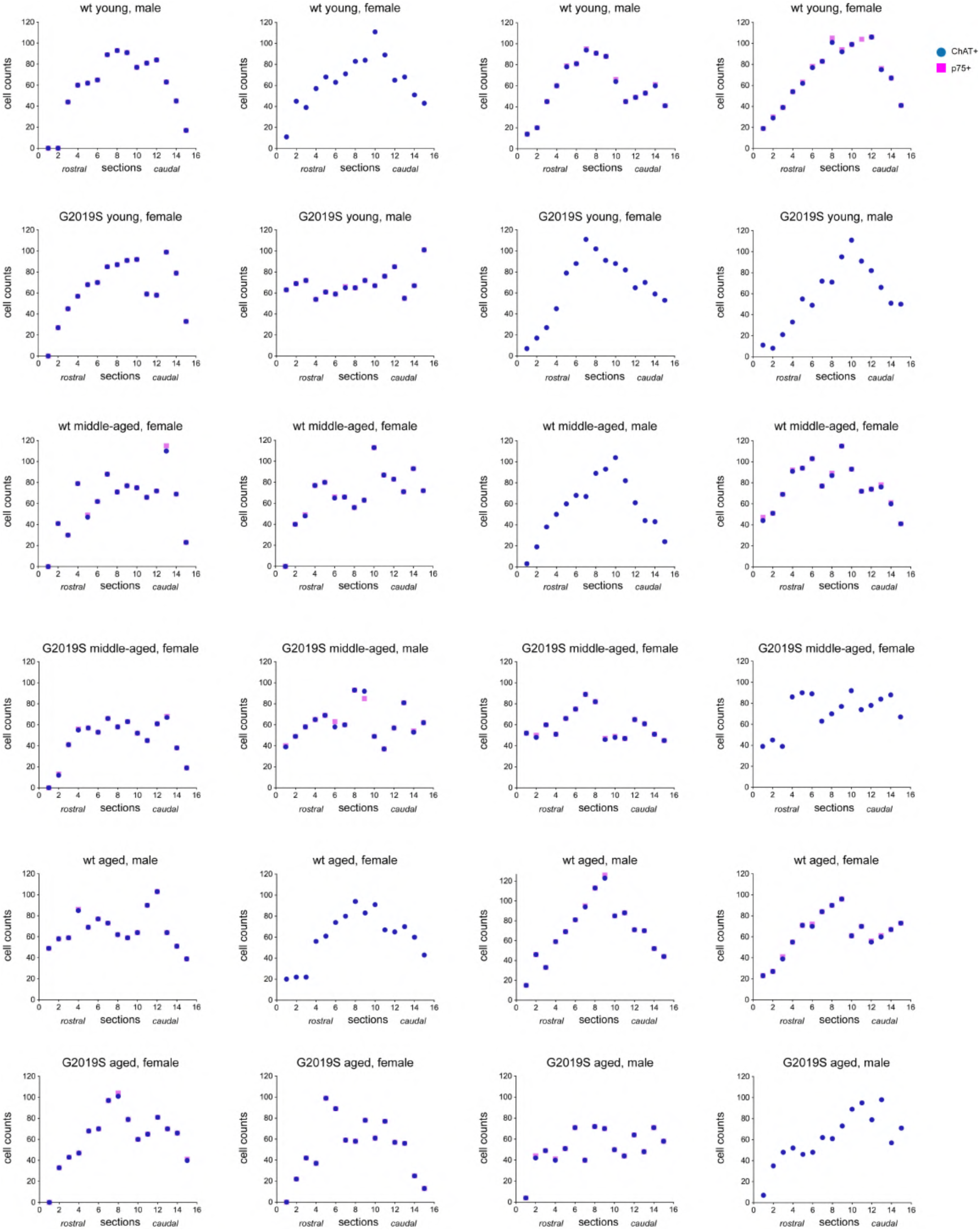
Cholinergic cell counts in MS. ChaT+ and p75+ cell counts in alternate sections through the medial septum from young adult (4-5 months), middle-aged (13-14 months) and aged (18-24 months) male and female mice as indicated.

**Figure 10 – Figure supplement 1.**
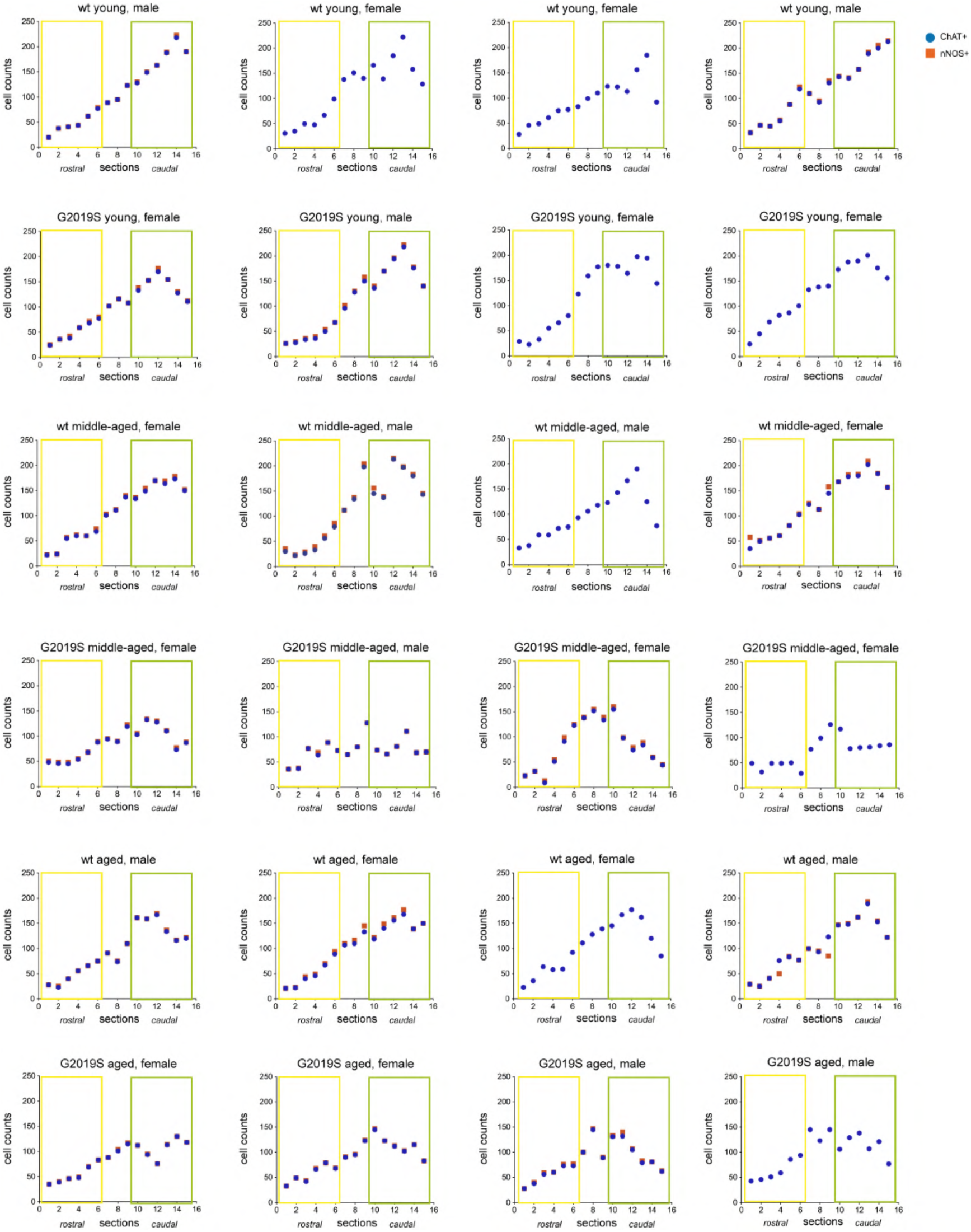
Cholinergic cell counts in PPN. ChaT+ and nNOS+ cell counts in alternate sections through the PPN from young adult (4-5 months), middle-aged (13-14 months) and aged (18-24 months) male and female mice as indicated.

## Notes

### Competing Interest Statement

The authors have declared no competing interest.

